# Selective YAP activation in Procr cells is essential for ovarian stem/progenitor expansion and epithelium repair

**DOI:** 10.1101/2021.11.17.468967

**Authors:** Jingqiang Wang, Lingli He, Zhiyao Xie, Wentao Yu, Lanyue Bai, Zuoyun Wang, Yi Lu, Chunye Liu, Junfen Fu, Lei Zhang, Yi Arial Zeng

## Abstract

Ovarian surface epithelium (OSE) undergoes recurring ovulatory rupture and OSE stem cells rapidly generate new cells for the repair. How the stem cell senses the rupture and promptly turns on proliferation is unclear. Our previous study has identified that Protein C Receptor (Procr) marks OSE progenitors. In this study, we observed decreased adherent junction and selective activation of YAP signaling in Procr progenitors at OSE rupture site. OSE repair is impeded upon deletion of Yap in these progenitors. Interestingly, Procr+ progenitors show lower expression of Vgll4, an antagonist of YAP signaling. Overexpression of Vgll4 in Procr+ cells hampers OSE repair and progenitor proliferation, indicating that selective low Vgll4 expression in Procr+ progenitors is critical for OSE repair. In addition, YAP activation promotes transcription of the OSE stemness gene *Procr*. The combination of increased cell division and Procr expression leads to expansion of Procr+ progenitors surrounding the rupture site. These results illustrate a YAP- dependent mechanism by which the stem/progenitor cells recognize the ovulatory rupture, and rapidly multiply their numbers, highlighting a YAP- induced stem cell expansion strategy.

## Introduction

During the adult reproductive cycles, the OSE undergoes recurring ovulatory rupture and repair^1, 2^. After ovulation, to maintain the physiological function and morphology of the ovary, the wound is completely closed within 12 hours to 3 days following rupture^3–5^. Cells surrounding the damaged sites are required to respond to the wound by turning on cell proliferation to supply sufficient cells as building block for regeneration^6^. Our previous study has identified that Procr+ OSE stem/progenitor cells are the major contributor for ovulatory rupture repair. Targeted ablation of these cells hampers the repair^7^. Interestingly, we observed that Procr+ cells expand instantly upon ovulation, reminiscent of a result of symmetric division^7^. It remains unknown how the stem cell senses the ovulation event, and what is the signal that triggers the instant stem cell expansion at the rupture site.

The cue for this stem/progenitor cell amplification likely comes from a particular extracellular signal occurring upon ovulation. One possibility is that the follicular fluid expelled during ovulation consists of Wnts and other potential niche signals^8–11^, which may regulate Procr+ stem/progenitor cell expansion. Another possibility is the involvement of mechanical force-induced signals during ovulation, resulting in Procr+ stem/progenitor cell expansion.

YAP (Yes-associated protein, also known as YAP1) signaling is an evolutionarily conserved pathway and a master regulator of organ size and tissue growth during animal development^12^. As a downstream effector, YAP is critical for regeneration in different organs, through triggering cell proliferation, cell survival or expansion of stem and progenitor cell compartments^13–19^. YAP is a transcriptional coactivator protein that shuttles between the cytoplasm and nucleus, and regulate expression of target genes, such as *Cyr61* and *Ctgf*, through binding with TEAD transcription factors^20–24^. Vgll4, a member of Vestigial-like proteins, serves as a transcriptional repressor of YAP through direct interactions with TEADs^25^. Previous studies from us and others have demonstrated the important roles of Vgll4 plays during development and regeneration in various tissues^26–29^. Cell-cell junctions links cells to each other in epithelial tissues, and is an upstream negative regulator of YAP^30, 31^. Mechanical forces regulate cell-cell adhesion stability, and cell-cell adhesion junctions may be intrinsically weak at high forces^32^. It has been shown that disruption of adherent junctions turns on YAP nuclear activities in lung stem/progenitor cells^33^. However, whether YAP signaling is implicated in ovulatory rupture repair is unknown.

In this study, we investigated how OSE stem/progenitor cells sense the rupture post ovulation and divide subsequently. We found that, in the proximity of rupture site, decreased adherent junction is associated with Yap nuclear localization in all cells, and conditional deletion of Yap in Procr+ cells hampers OSE repair. Interestingly, only Procr+ OSE cells displayed a low level of Vgll4, allowing YAP signaling activation. We generated a new *tetO- Vgll4* mouse. Ectopic expression of Vgll4 in the stem/progenitor cells using *Procr-rtTA;tetO-Vgll4* mice blocked OSE ovulatory repair. Moreover, we found that YAP signaling activation resulted in Procr+ cells expansion at the rupture site, through the combination of inducing cell division, and directly activating Procr transcription. The activation of Procr is essential, as when Procr was deleted, stemness property was lost and OSE repair was hindered.

## Results

### Decreased E-Cadherin expression at the rupture site and selective activation of YAP signaling in Procr+ cells

To investigate what could be the potential extracellular stimuli at the rupture site, we performed immunostaining of various adherent or tight junction components on ovary sections. To increase rupture incidences, superovulation was induced by injection of PMSG and HCG, and the ovaries were harvested at 0.5 days after HCG injection, when ovulation just occurred (Fig. S1a). Interestingly, we found that E-Cadherin staining is markedly decreased at the proximal region of rupture (defined as within 20 cells on one side of the rupture in section) compared to other regions, i.e., rupture distal region (Fig. 1a) and non-rupture region (Fig. S1b). As adherent junction has been implicated as a modulator of YAP signaling^31, 34–36^, we examined YAP activities at the rupture area by immunostaining. We observed increased incidence of nuclear YAP at the proximal region of rupture compared to other regions (Fig. 1b-c, Fig. S1c). These results suggest that compromised adherent junction resulted from ovulatory rupture could induce YAP nuclear localization in OSE cells surrounding the wound.

**Fig 1.**
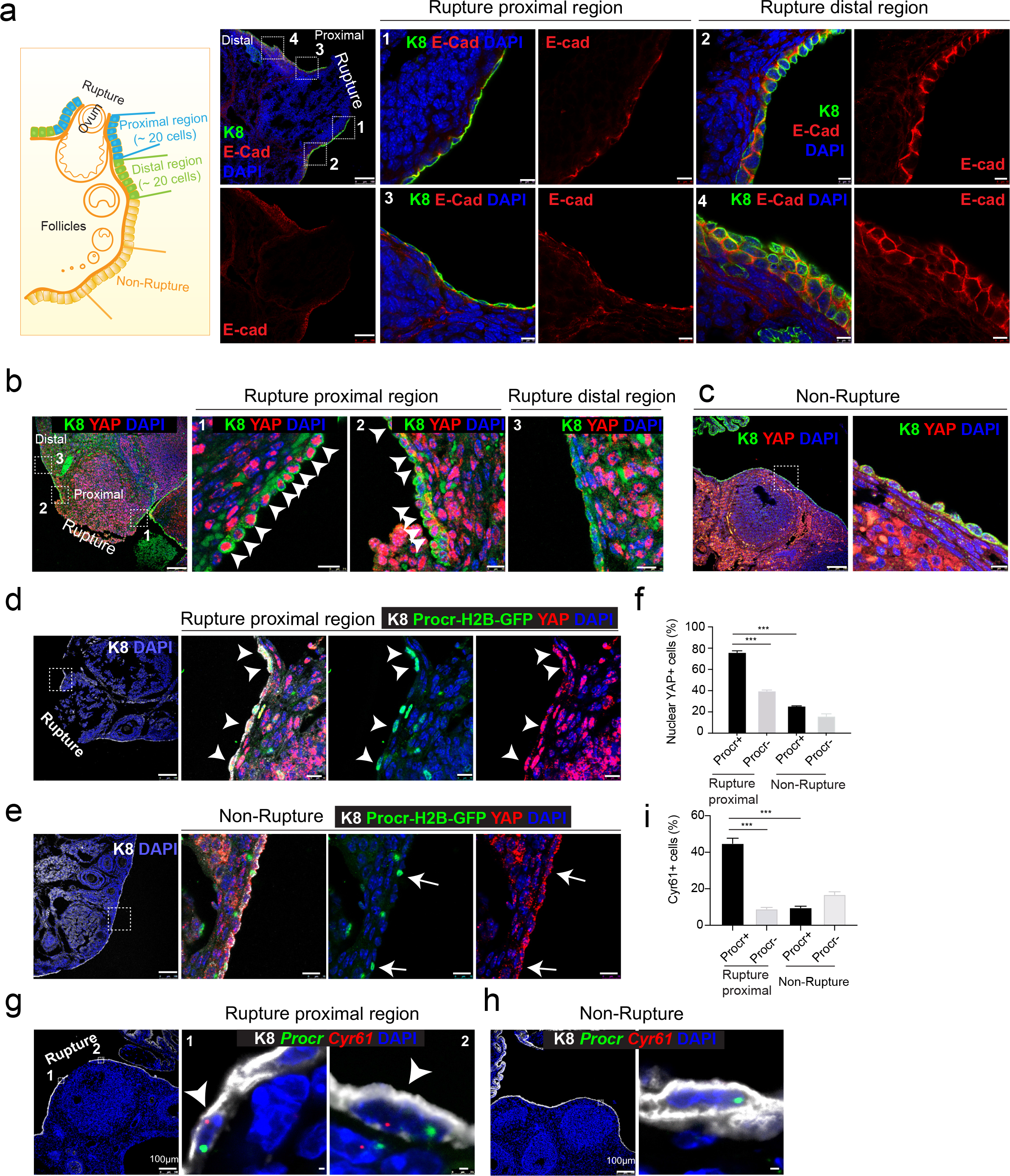
Rupture-induced YAP signaling activation is preferentially activated in Procr+ progenitors at the rupture sites. a, Sections from wildtype ovaries at ovulation stage were stained with Krt8 (K8) and E-cadherin (E-Cad). Confocal images showed less E-cad in the OSE of proximal regions surrounding the rupture sites (view #1, #3 in a) compared with distal regions (view #2, #4 in a). Scale bar, 100μm for zoom out and 10μm for zoom in. n=3 mice and 15 images. b-c, Sections from wildtype ovaries at ovulation stage were stained with K8 and YAP. Confocal images showed YAP nuclear localization in the OSE was only observed in the proximal regions surrounding the rupture sites (b), but not in the distal regions (b) or the non-rupture sites (c). Scale bar, 100μm for zoom out and 20μm for zoom in. n=3 mice and 15 images. d-f, *Procr-rtTA;TetO-H2B-GFP^+/-^* mice were fed with doxycycline for 3 days and harvested at ovulation stage. Confocal images of ovarian sections with Krt8 (K8) and YAP staining (d-e) and quantification (f) were showed. Nuclear YAP staining is preferentially detected in Procr+ (histone 2B-GFP+) cells in rupture proximal region (arrowheads in d), whereas at the non-rupture site, YAP staining was cytoplasmic regardless in Procr+ (arrows in e) or Procr- cells (e). Scale bar, 100μm for zoom out and 10μm for zoom in. n=3 mice and 15 images. One-way ANOVA with Tukey test is used for comparison of multiple groups. ***P<0.001. g-i, Combination of *Procr* and *Cyr61* double fluorescent *in situ* with K8 antibody immunohistochemistry staining (g-i). Confocal images showed co-localization of *Procr* and *Cyr61* in the OSE at the rupture sites (arrowhead in g), while at non-rupture regions, both Procr+ and Procr- cells had low incidence of *Cyr61* expression (h). Quantification showed increased *Cyr61* expression in Procr+ cells at rupture sites compared with Procr- cells at rupture sites or Procr+ cells at non-rupture regions (i). Scale bar, 100μm for zoom out and 1μm for zoom in. n=3 mice and 15 images. One-way ANOVA with Tukey test is used for comparison of multiple groups. ***P<0.001.

Our previous study has established that Procr+ progenitor cells surrounding the wound instantly proliferate upon rupture and are responsible for OSE repair^7^. We therefore investigated whether Procr+ cells close to the rupture site are associated with YAP signaling activities. We performed YAP immunostaining using *Procr-rtTA*;*tetO-H2B-GFP* reporter, in which H2B-GFP signal marks Procr-expressing cells. Superovulation was induced in these animals by PMSG and HCG injections, and ovaries were harvested 0.5 days after HCG injection. We found that Procr+ (H2B-GFP+) cells at the rupture proximal region (referred to as rupture site from here on) have significantly higher nuclear YAP staining (75.9±1.7%) compared to Procr- cells (39.6±1.0%) (Fig. 1d, 1f), or compared to Procr+ cells at the non-rupture region (Fig. 1e-f). This was further validated by RNA double *in situ* hybridization with *Procr* and a YAP target gene *Cyr61*. We found that, at the rupture site, *Cyr61* is preferentially activated in Procr+ OSE cells, with 44.8±2.9% of Procr+ cells being Cyr61+, which is markedly higher than that of Procr- cells (8.9±0.8%) (Fig. 1g, 1i). At the non-rupture site, both Procr+ and

Procr- cells had rather low expression of *Cyr61* expression (Fig. 1h-i). Together, these results suggest that YAP signaling was specifically activated in Procr+ cells at the rupture site. Considering the role of YAP signaling in promoting cell proliferation, there results are in line with our previous observations that only Procr+, but not Procr-, cells at the rupture site displayed increased proliferation^7^.

### Deletion of Yap in Procr+ cells hinders rupture repair and progenitor proliferation

To investigate whether YAP signaling is important for OSE repair, we deleted YAP specifically in Procr+ cells using *Procr-CreER*;*Yap^fl/fl^* mice (Yap- cKO). *Yap^fl/fl^* mice was used as control (Ctrl). Tamoxifen (TAM) was administered in 4-week-old mice, following by superovulation at 2 days after TAM injection (Fig. 2a). The impact on OSE repair by Yap deletion was analyzed by ovary whole-mount imaging. At 4.5d pi (ovulation), the two groups had similar ruptures (Fig. 2b, 2e). At 6d pi, Ctrl ovaries underwent rapid repairing (Fig. 2c), and the OSE was completely recovered by 7.5d pi (Fig. 2d). In contrast, the OSE repair in Yap-cKO ovaries was significantly delayed at both 6d pi and 7.5d pi (Fig. 2c-e). The efficacy of Yap deletion and the reduced expression of the target gene *Cyr61* in OSE cells were validated by qPCR analyses (Fig. S1d).

**Fig 2.**
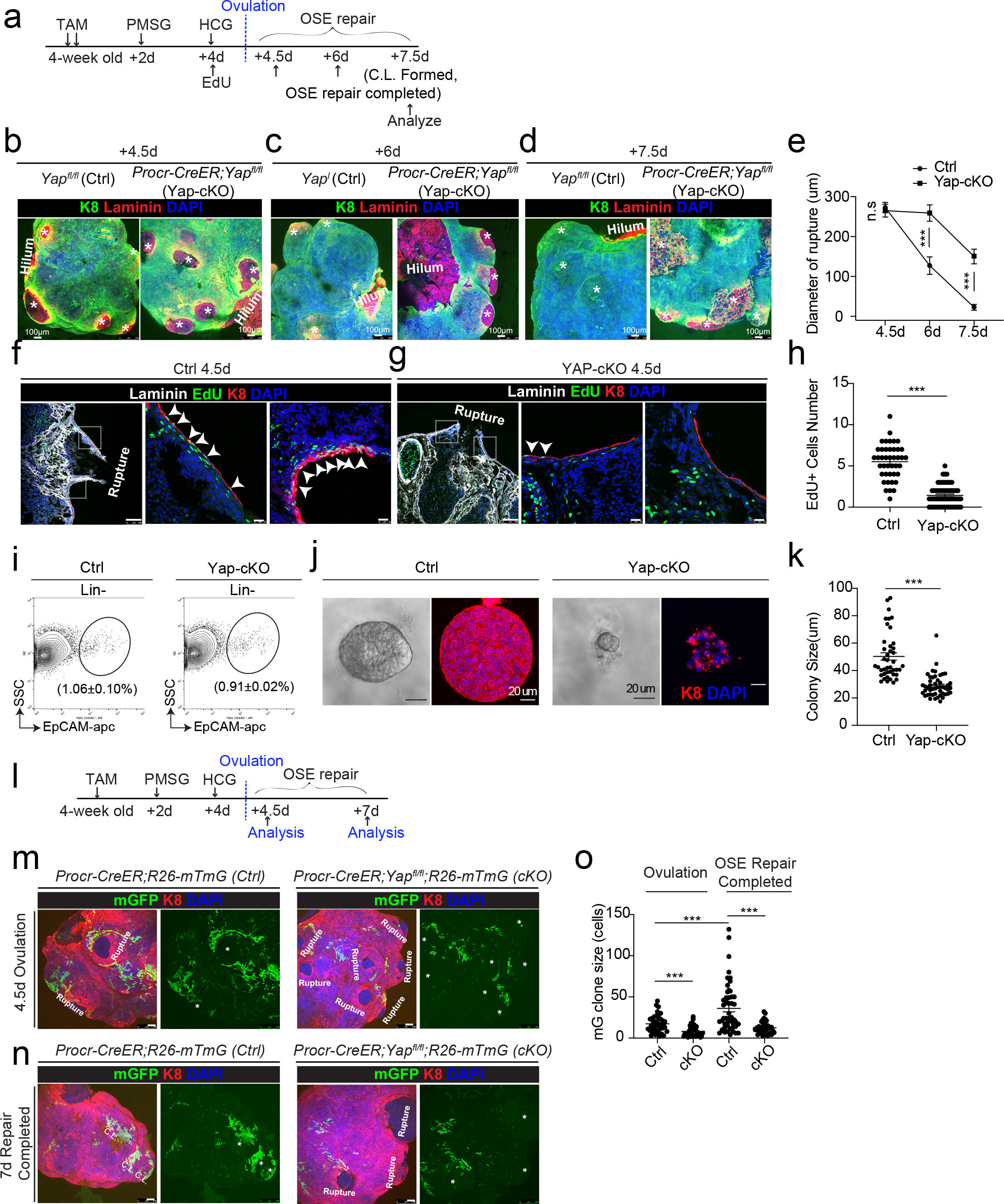
Deletion of YAP in Procr+ cells hinders OSE rupture repair and progenitor proliferation. a, Illustration of TAM induction and superovulation strategy. b-e, Yap was deleted in Procr+ cells using *Procr-CreER;Yap^fl/fl^* mice (Yap-cKO), and *YAP^fl/fl^* mice was used as control (Ctrl). Ovary whole-mount staining with K8 and Laminin was performed (b-d) and the wound size in diameter was quantified (e). At 4.5d (ovulation) (star) Ctrl and Yap-cKO ovaries had comparable wound size (* in b, e). At 6d (OSE repair ongoing), the wounds in Ctrl ovary were significantly smaller than those in Yap-cKO ovary (c, e). At 7.5d (repair completed), the wound was completely repaired in Ctrl, while the Yap- cKO ovary still showed obvious wounds (star) (* in d, e). Scale bar, 100μm. n=3 pairs of mice. f-h, Ctrl and Yap-cKO mice were subjected to 12hrs EdU incorporation and were harvested at 4.5 days (Ovulation stage). Representative images (f-g) and quantification (h) were showed. Out of 20 cells next to the rupture on one side, the numbers of EdU+ cells (arrowhead) in the OSE of rupture site decreased from 5.52±0.33 cells in Ctrl to 1.42±0.16 cells in Yap-cKO. Scale bar, 100μm for zoom out and 20μm for zoom in. n=3 pairs of mice. Unpaired two-tailed t test is used for comparison. ***P<0.001. i-k, Total OSE cells from *Procr-CreER;Yap^fl/fl^* mice (Yap-cKO), and *Yap^fl/fl^* mice (Ctrl) were isolated by FACS at 4.5 days (Ovulation stage) (i), followed by culture in 3D Matrigel for 7 days. Representative bright-field and confocal images of K8 staining were shown (j). Colony sizes in diameter were measured (k). Scale bar, 20μm. Data are pooled from three independent experiments and displayed as mean±s.e.m. Unpaired two-tailed t test is used for comparison. ***P < 0.001. l, Illustration of lineage tracing, deletion of YAP and superovulation strategy. m-o, *Procr-CreER;YAP^fl/fl^;R26-mTmG* (Yap-cKO) and *Procr-CreER;R26- mTmG* (Ctrl) mice were used. At 4.5pi (ovulation), ovary whole-mount confocal imaging showed zones of concentrated GFP+ cells surrounding the rupture site in Ctrl, while fewer GFP+ cells were seen in Yap-cKO ovary (m). At 7pi (repair completed), ovary whole-mount confocal imaging showed large GFP+ patches located at corpus luteum (C.L.) in Ctrl, while rare GFP+ cells surrounding the unrepaired wound in Yap-cKO ovary (n). Quantification showed significantly fewer GFP+ cells in Yap-cKO compared with Ctrl in both ovulation stage and repair completed stage. Quantification showed an expansion of GFP+ cell numbers in Ctrl mice during the tracing and no expansion in Yap-cKO (o). Scale bar, 100μm. n=3 pairs of mice. ***P < 0.001.

To analyze the proliferative capacity of Procr+ OSE cells, mice were subjected to 12 h of EdU incorporation before harvesting the ovaries (Fig. 2a). When analyzed at 4.5d pi (ovulation), the number of proliferating OSE cells at rupture site (defined as 20 cells on one side from the opening) were significantly decreased from 5.5±0.3 EdU+ in Ctrl to 1.4±0.2 EdU+ in Yap-cKO (Fig. 2f-h). The impact to cell proliferation was further analyzed *in vitro.* Our previous study has established that Procr+, but not Procr-, OSE cells can form colonies *in vitro*^7^. At 4.5d pi, total OSE cells were isolated from both Ctrl and Yap-cKO mice (Fig. 2i), and placed in culture as previously described^7^. Deletion of Yap in Procr+ cells drastically inhibited OSE colony formation (Fig. 2j-k).

To visualize the contribution of Procr+ progenitors toward the repair in the presence or absence of Yap, we performed *in vivo* lineage tracing. TAM was administered to 4-week-old mice to simultaneously delete Yap and initiate lineage tracing in Procr+ cells (Fig. 2l). At 4.5d pi, control (*Procr-CreER*;*R26- mTmG*) ovary displayed a zone of mGFP+ cells that are the progeny of

Procr+ progenitors surrounding the rupture sites (Fig. 2m). In contrast, Yap- cKO (*Procr-CreER*;*Yap^fl/fl^;R26-mTmG*) ovaries have markedly fewer mGFP+ cells around the wound (Fig. 2m, 2o), supporting the notion that the activity of Procr+ progenitors was hampered at the beginning of the repairing process. At 7d pi, control ovaries had generated patches of mGFP+ cells covering the newly formed corpus luteum (Fig. 2n). However, Yap-cKO ovaries still had obvious openings with few mGFP+ cells (Fig. 2n-o). Together, these results suggest that YAP signaling activation is crucial for the proliferation of Procr+ progenitor cells and the timely repair of OSE after rupture.

### An intrinsic lower level of Vgll4 in Procr+ cells is essential for their progenitor property and OSE rupture repair

Next, we investigated what could be the reason that YAP signaling is specifically activated in Procr+ cells. Vgll4 is a negative regulator of YAP by inhibiting the binding of YAP and TEAD4^26^. We FACS-isolated Procr+ cells and Procr- cells from the rupture sites (Fig. 3a). qPCR analysis indicated that Procr+ cells have lower level of *Vgll4* compared to Procr- cells (Fig. 3b). This was further validated by Vgll4 immunostaining using *Procr-rtTA*;*tetO-H2B- GFP* reporter, in which H2B-GFP signals mark Procr-expressing cells. Consistent with the qPCR results, Procr+ cells also exhibited lower Vgll4 protein expression compared to Procr- cells (Fig. 3c-e).

**Fig 3.**
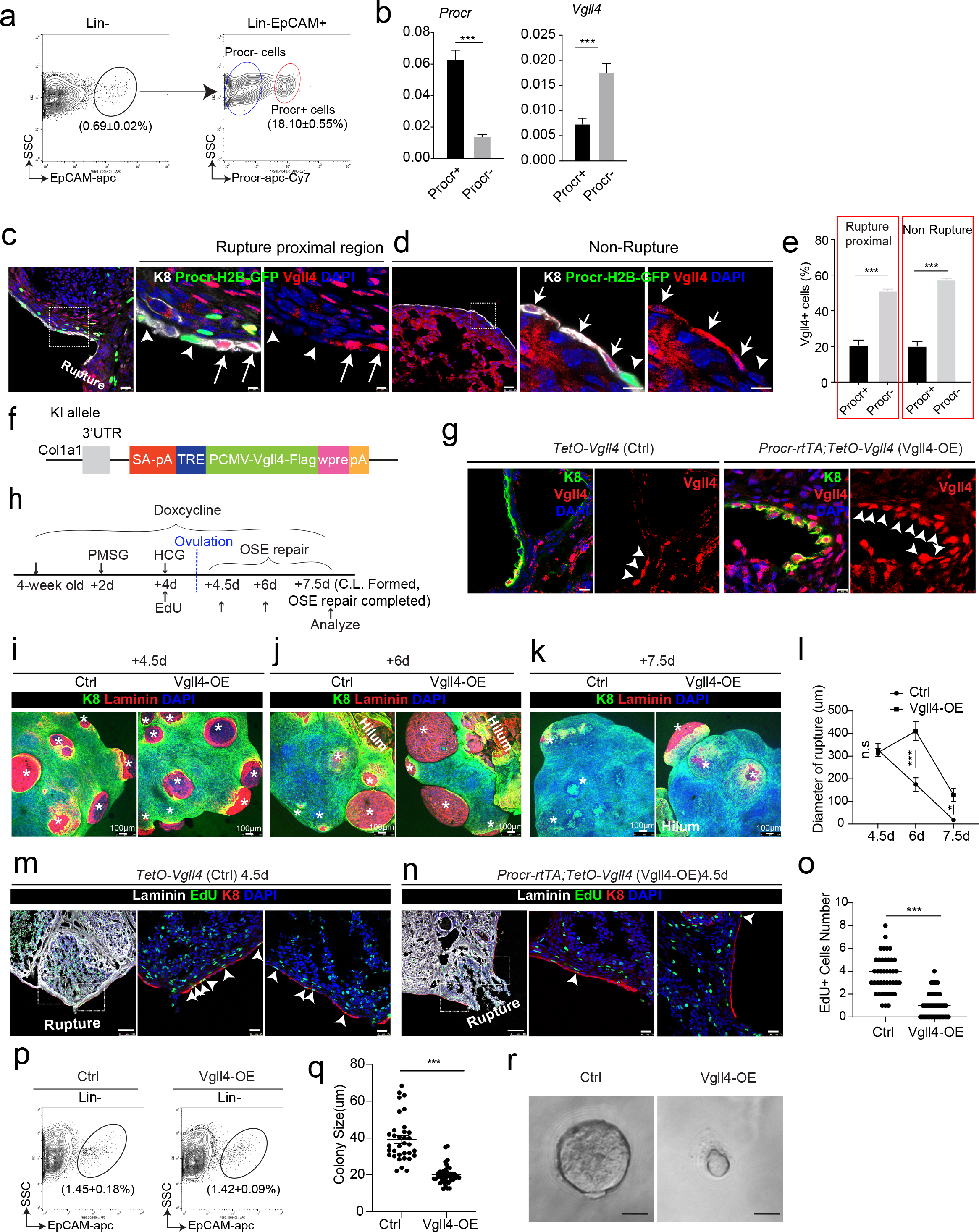
An intrinsic lower level of Vgll4 in Procr+ cells is essential for Procr+ cells’ stemness and OSE rupture repair. a-b, At ovulation stage, Procr+ and Procr- OSE cells (Lin-, EpCAM+) were FACS-isolated (a). qPCR analysis showed the lower *Vgll4* level in Procr+ cells (b). Data are pooled from 3 independent experiments and presented as mean±s.e.m. ***P<0.001. c-e, *Procr-rtTA;TetO-H2B-GFP* mice were administered with PMSG and HCG to induce superovulation, and fed with doxycycline for 3 days before harvest. Ovarian sections were stained with Vgll4 and K8. Representative images showed that at both rupture proximal region (c) and non-rupture region (d), H2B-GFP- (Procr-) OSE cells have high Vgll4 expression (arrows in c, d), while H2B-GFP+ (Procr+) OSE cells have no Vgll4 expression (arrowheads in c, d). Scale bar, 20μm for zoom out and 5μm for zoom in. Quantification of the staining was shown in (e). n=3 mice and 15 images. Unpaired two-tailed t test is used for comparison. ***P<0.001. f-g, Targeting strategy and validation of *TetO-Vgll4* knock-in mouse (f-g). A cassette of TetO-Vgll4-Flag-wpre-polyA was knocked in behind 3’UTR of *Col1a1* gene (f). Immunohistochemistry staining of Vgll4 in the ovaries indicated more Vgll4+ OSE cells at the rupture sites (g). Scale bar, 10μm. n=3 pairs of mice. h-l, Illustration of superovulation and overexpression of Vgll4 in Procr+ cells (h). Ovary whole-mount confocal images of K8 and Laminin showed that at 4.5d (ovulation), Ctrl (*TetO-Vgll4*) and Vgll4-OE (*Procr-rtTA;TetO-Vgll4*) ovaries have similar wound sizes (* in i) . At 6d (repair ongoing), the wound sizes in Ctrl mice were smaller than those in Vgll4-OE (* in j). At 7.5d (repair completed), Ctrl ovary had completely repaired, while Vgll4-OE mice had obvious opening (* in k). Scale bar, 100μm. The sizes of the wound in diameter were quantified (l). n=3 pairs of mice. m-o, The mice were harvested at 4.5 days (ovulation) after 12hrs EdU incorporation. Representative images (m, n), and quantification (o) showed the number of EdU+ cells in the OSE surrounding the rupture site (arrowheads in m) decreased from 3.73±0.26 in Ctrl to 1.04±0.15 in Vgll4-OE (arrowheads in n). Scale bar, 100μm for zoom out and 20μm for zoom in. n=3 pairs of mice. Unpaired two-tailed t test is used for comparison. ***P<0.001. p-r, Total OSE cells were isolated by FACS from Ctrl and Vgll4-OE at 4.5 days (ovulation) (p), and cultured in 3D Matrigel. At day 7 in culture, colony sizes were measured in diameter (q), and representative images were shown (r) out of 15 images in each group. Scale bar, 20μm. Data are pooled from three independent experiments and displayed as mean±s.e.m. Unpaired two-tailed t test is used for comparison. ***P < 0.001.

To examine whether the reduced level of Vgll4 is significant for the selective YAP signaling activation in Procr+ cells and rupture repair, we set to overexpress Vgll4 specifically in Procr+ cells. A new *tetO-Vgll4* mouse line was generated, by inserting a tetO-Vgll4-Flag-wpre-polyA cassette behind the 3’UTR of the *Col1a1* gene (Fig. 3f and Fig. S2a-c). Subsequently, *Procr- rtTA*;*tetO-Vgll4* (Vgll4-OE) mice were generated by genetic crosses with *TetO- Vgll4* as control (Ctrl). The efficacy of overexpression was validated by western blotting and qPCR, showing increased expression of Vgll4 and decreased expression of *Cyr61* in Vgll4-OE cells (Fig. S2d-e). Furthermore, immunostaining confirmed the increased number of Vgll4-high cells in the OSE layer of Vgll4-OE mice (Fig. 3g). For this experiment, superovulation was performed to 4-week-old mice and DOX was fed throughout the process (Fig. 3h). The impact of Vgll4 overexpression was analyzed throughout the repairing process, at 4.5d pi (ovulation), 6d pi (OSE repair ongoing) and 7.5d pi (OSE repair completed) by ovary whole-mount imaging. We found that the rupture in Ctrl and Vgll4-OE ovaries are comparable at 4.5d pi (Fig. 3i, 3l). At 6d pi, while Ctrl ovaries had sights of wound closure, Vgll4-OE ovaries still showed larger areas of rupture (Fig. 3j, 3l). At 7.5d pi, Ctrl ovaries displayed complete OSE, whereas the repair in Vgll4-OE ovaries was obviously delayed (Fig. 3k-l).

Next, we examined whether overexpression of Vgll4 affects progenitor proliferation. At 4.5d pi (ovulation), the number of proliferating OSE cells at rupture site were significantly decreased from 3.7±0.3 in Ctrl to 1.0±0.2 in Vgll4-OE (Fig. 3m-o). At 4.5d pi, total OSE cells were isolated and cultured *in vitro* for 7 days (Fig. 3p). Consistently, overexpression of Vgll4 inhibits cell proliferation and colony formation (Fig. 3q-r). Together, these results suggest that overexpression of Vgll4 in Procr+ cell impaired Procr+ cell proliferation and ovulatory rupture repair.

### YAP signaling promotes Procr+ cells expansion at rupture site

We have previously found that Procr+ progenitor cells expand instantly at the periphery of the rupture site upon ovulation^7^. To investigate whether YAP signaling activation is linked to the expansion of Procr+ progenitor cells, TAM was administered to 4-week-old *Procr-CreER*;*Yap^fl/fl^* (Yap-cKO) and *Yap^fl/fl^* (Ctrl) mice for 2 times, followed by superovulation. At 4.5d pi (ovulation), FACS analysis showed a dramatic decrease of Procr+ progenitor population when Y*ap* was deleted (Fig. 4a-c).

**Fig 4.**
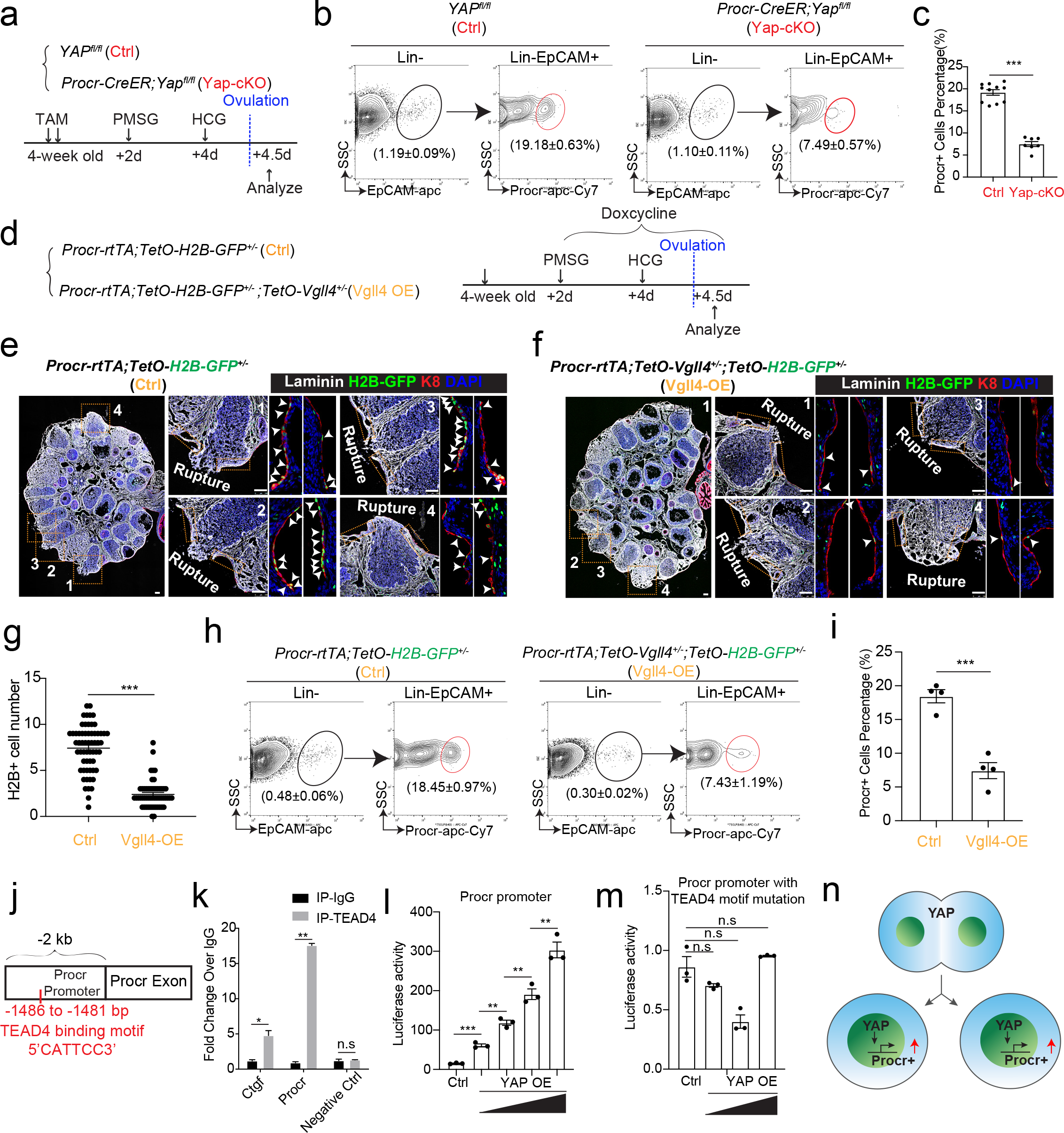
YAP signaling promotes Procr+ cells expansion at rupture sites through a combination of promoting cell division and enhancing Procr expression. a-c, Illustration of superovulation and analysis strategy as indicated using *YAP^fl/fl^* (Ctrl) and *Procr-CreER;YAP^fl/fl^* (Yap-cKO) mice (a). At ovulation stage, the percentage of Procr+ OSE cells in Ctrl and Yap-cKO were FACS analyzed (b) and quantified (c). n = at least 3 mice in each group and displayed as mean±s.e.m. Unpaired two-tailed t test is used for comparison. ***P < 0.001. d-i, Illustration of superovulation and analysis strategy as indicated using *Procr- rtTA;TetO-H2B-GFP^+/-^* (Ctrl) and *Procr-rtTA;TetO-H2B-GFP^+/-^;TetO-Vgll4^+/-^* (Vgll4-OE) mice (d). At ovulation stage, ovary section imaging showed that at the rupture sites, the number of H2B-GFP+ (Procr+) cells in Ctrl (arrowheads in e) are higher than those in Vgll4-OE (arrowheads in f). Scale bar, 100μm. Quantification was shown in (g). n=3 pairs of mice and 15 images in each group. Unpaired two-tailed t test is used for comparison. ***P<0.001. The percentage of Procr+ OSE cells were analyzed by FACS at ovulation stage (h). The percentage of Procr+ cells in Ctrl are higher than that in Vgll4-OE (h, i). n = at least 3 mice and displayed as mean±s.e.m. Unpaired two-tailed t test is used for comparison. ***P < 0.001. j-k, Illustration of Tead4 motif in Procr promoter region (j). TEAD4 ChIP-qPCR analysis using cultured primary OSE cells showed the enrichment of Procr promoter, and Ctgf promoter was used as positive control (k). n=2 biological repeats. Unpaired two-tailed t test is used for comparison. **P<0.01, *P<0.05, n.s, not significant. l-m, Analysis of luciferase reporter activity driven by WT (l) and Tead4 motif (- 1486 to-1481bp) deleted- Procr promoter (m) in HEK193T cells transfected with increased amount of YAP overexpression plasmids. Data are pooled from three independent experiments and displayed as mean±s.e.m. Unpaired two-tailed t test is used for comparison. ***P < 0.001, **P < 0.01, n.s, not significant. n, A proposed model of which YAP signaling promotes Procr+ cells expansion at rupture site through a combination of promoting cell division and enhancing Procr expression.

To better visualize the change of Procr+ progenitor cells under the influence of Yap signaling, we generated *Procr-rtTA*;*TetO-H2B-GFP^+/-^*;*TetO- Vgll4^+/-^* mice (Vgll4-OE). Superovulation was performed to 4-week-old mice and DOX was fed throughout the experiments to maintain the expression of H2B-GFP in Procr+ cells (Fig. 4d). When analyzed at 4.5d pi (ovulation), at the wound edge (defined as 20 cells on one side from the opening) of control ovary (*Procr-rtTA*;*TetO-H2B-GFP^+/-^*), there were about 7.4±0.3 H2B-GFP+ cells expressing the peak level of GFP (Fig. 4e, 4g). In contrast, in Vgll4-OE ovary (*Procr-rtTA*;*TetO-Vgll4;TetO-H2B-GFP^+/-^*), only 2.4±0.2 H2B-GFP+ cells were observed at the wound edge (Fig. 4f-g). FACS analysis also showed that the percentage of Procr+ progenitor population decreased significantly from 18.5±1.0% in Ctrl to 7.4±1.2% in Vgll4-OE at ovulation stage (Fig. 4h-i).

The proliferative activity of Procr+ cells was further evaluated *in vitro*. We isolated OSE cells from control (*Procr-rtTA*;*TetO-H2B-GFP^+/-^*) and Vgll4-OE (*Procr-rtTA*;*TetO-Vgll4*;*TetO-H2B-GFP^+/-^*) mice and placed in culture, followed by live imaging to document the division of H2B-GFP+ (Procr+) cells (Fig. S3a-b). In control cells, we observed frequent division of Procr+ cells, and in most cases, it was one Procr+ cell dividing into two Procr+ cells (Fig. S3a). But in Vgll4-OE, we could hardly observe cell division (Fig. S3b). Together, these results suggest that inhibition of YAP signaling, by either Yap-deletion or Vgll4- OE, impairs the expansion of Procr+ progenitors upon rupture.

### YAP signaling enhances Procr expression

It is unclear how YAP maintains Procr expression during or after cell division. Thus, we investigated the association of YAP activation and Procr expression. OSE cells were isolated from *Procr-rtTA*;*TetO-H2B-GFP^+/-^* mice, and cultured on glass (YAP activation) or soft condition (0.48kPa, YAP inactivation) (Fig. S4a-b). DOX was added 2 days before harvest. Consistent with the notion, we found that, in soft condition, YAP was mostly cytoplasmic and most OSE cells are H2B-GFP- (Fig. S4a). In contrast, most OSE cells are H2B-GFP+ in stiff condition and YAP was found in the nucleus (Fig. S4b). These observations suggest that YAP activation might induce Procr expression. We verified by qPCR that *Procr* expression is upregulated in stiff conditions (Fig. S4c). Our results support the notion that YAP activation induces Procr expression.

To further investigate whether YAP regulates Procr expression, we knocked down YAP by shRNA in OSE culture and found that this inhibits Procr expression (Fig. S4d-e). Furthermore, blocking YAP activation by Verteporfin (VP) or Vgll4 overexpression also resulted in lower Procr expression (Fig. S4f-g). These results suggest that inhibiting YAP signaling suppresses Procr expression.

To investigate whether YAP/TEAD4 directly regulate Procr expression, we analyzed the promoter of *Procr*. A Tead4 binding motif (5’-CATTCC-3’) was found at the proximal promoter of *Procr* (−1486 to -1481 bp) (Fig. 4j). ChIP- qPCR showed that Tead4 could directly bind to the Procr promoter (Fig. 4k). Therefore, we examined whether this Tead4 binding motif is responsible for induction of Procr expression by Yap. While Yap induced the wild type promoter-luciferase in a dose-dependent manner (Fig. 4l), it could not activate the mutant reporter with the deletion of the Tead4 binding motif (Fig. 4m).

These results suggest that YAP directly promotes Procr expression. Together, our data support a model that YAP signaling promotes expansion of Procr+ cells at rupture site through a combination of increased cell division and Procr expression (Fig. 4n).

### Procr is essential for the progenitor property

The up-regulation of Procr expression coupled with YAP-induced cell division implies that the expression of Procr may be important for keeping the stem cell property in OSE. To assess the significance of Procr, we utilized a *Procr-flox* allele (Liu and Zeng unpublished) and specifically deleted Procr in the progenitor using *Procr^CreER/fl^* (Procr-cKO) mice. TAM was administered in 4- week-old mice for two times, followed by superovulation at 2 days after TAM injection (Fig. 5a), and the phenotype was analyzed by ovary whole-mount imaging. Ctrl and Procr-cKO ovaries formed comparable ruptures at 4.5d pi (ovulation) (Fig. 5b, 5e). At 6d pi (OSE repair ongoing), Procr-cKO ovaries showed bigger openings compared to control (Fig. 5c, 5e). At 7d pi (OSE repair completed), Ctrl ovaries were covered by complete OSE, whereas Procr cKO ovaries still had regions with unrepaired OSE (Fig. 5d-e). Furthermore, at 4.5d pi (ovulation), the ovaries were harvested after 12 h of EdU incorporation. The number of proliferated OSE cells at rupture site decreased from 5.5±0.3 cells in control (*Procr^fl/+^*) to 1.4±0.2 cells in Procr-cKO (Fig. 5f-h). After deletion of Procr *in vivo*, total OSE cells were isolated and cultured *in vitro* for 7 days (Fig. 5i). We found that deletion of Procr inhibits the proliferation of progenitor cells, resulting in reduced colony sizes (Fig. 5j-l). Overall, these data suggest that Procr is essential for progenitor property upon rupture.

**Fig 5.**
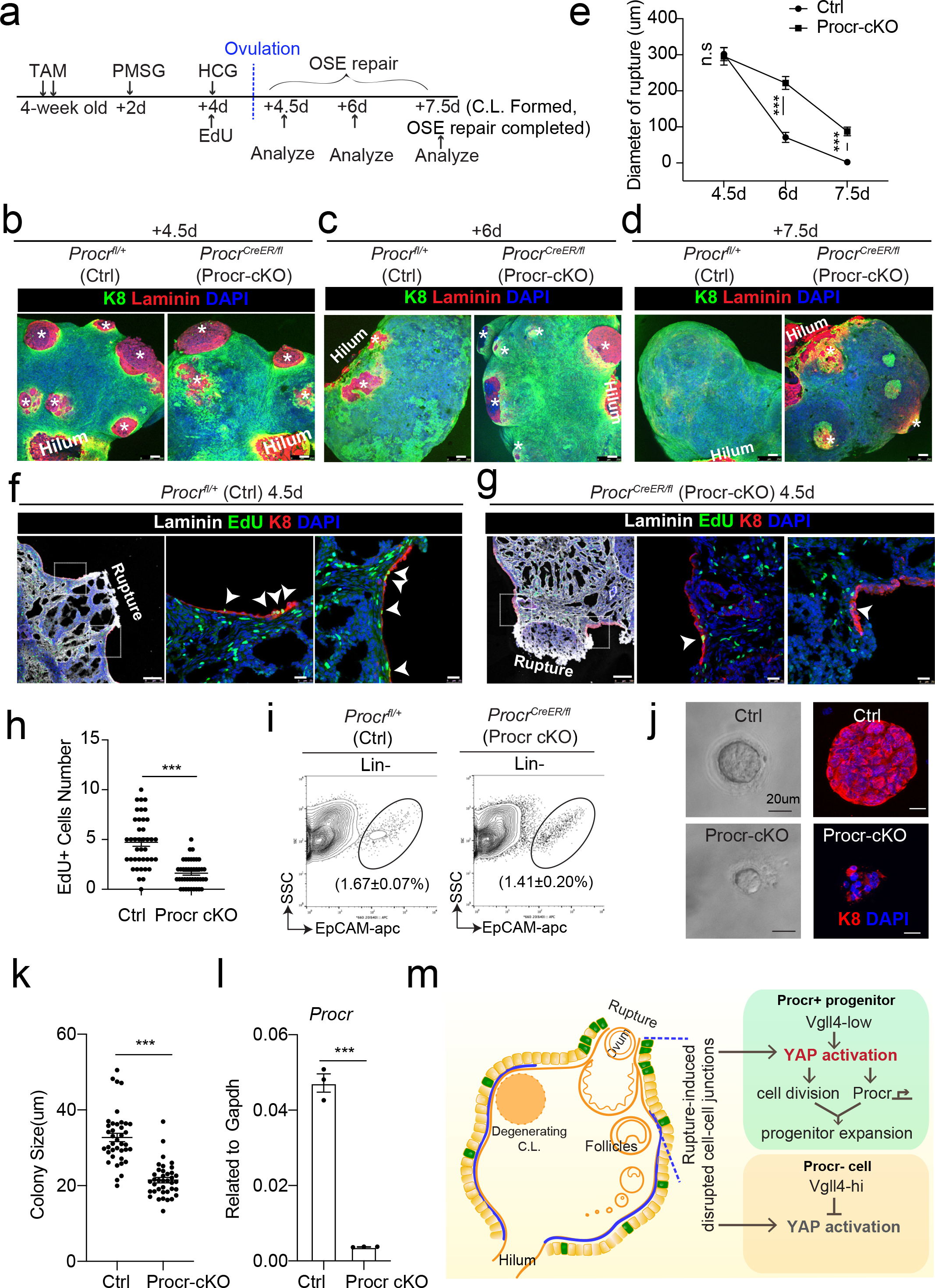
Procr is essential for the progenitor property. a-e, Illustration of superovulation and deletion of *Procr* in Procr+ cells using *Procr^CreER/fl^* mice (Procr-cKO), and *Procr^fl/+^* mice (Ctrl) (a). Ovary whole-mount confocal imaging of K8 and Laminin showed that at 4.5d (ovulation), Ctrl and Procr-cKO have similar wound sizes (* in b). At 6d (OSE repair ongoing), the wound sizes in Ctrl mice were smaller than those in Procr-cKO (* in c). At 7.5d (repair completed), Ctrl ovary had completely repaired, while Procr-cKO remained obvious opening (* in d). Scale bar, 100μm. Quantification of the wound size in diameter was shown in (e). n=3 pairs of mice. Unpaired two-tailed t test is used for comparison. ***P<0.001. n.s, not significant. f-h, Post 12hrs EdU incorporation, the mice were harvested at 4.5 days (ovulation) (a). Representative images showed the number of EdU+ cells (arrowhead) in the OSE surrounding the rupture site decreased from 4.73±0.40 in Ctrl (arrowheads in f) to 1.62±0.20 in Procr-cKO (arrowheads in g). Scale bar, 100μm for zoom out and 20μm for zoom in. Quantification of was shown in (h). n=3 pairs of mice. Unpaired two-tailed t test is used for comparison. ***P<0.001. i-l, Total OSE cells from Ctrl and Procr-cKO were isolated by FACS (i), followed by culture in 3D Matrigel. At culture d7, representative bright-field and confocal images with K8 staining showed that OSE cells with Procr-cKO form markedly smaller colonies compared to Ctrl (j). Colony sizes were quantified in (k). qPCR analysis validated the deletion efficiency of *Procr* in OSE cells of Procr-cKO (l). Data are pooled from three independent experiments and displayed as mean±s.e.m. Unpaired two-tailed t test is used for comparison. ***P < 0.001. Scale bar, 20μm. m, A proposed model of YAP activation in Procr+ cells promoting OSE progenitor cell expansion. Procr+ OSE progenitors have intrinsically lower level of Vgll4 compared to Procr- OSE cells. At ovulation, cell-cell junctions at rupture site were disrupted, which induces the possibility of YAP activation in all OSE cells surrounding the rupture. However, the lower expression of Vgll4 in Procr+ cells allowed YAP activation in the progenitor cells at this area. YAP activation in Procr+ cells promoted cell division, and importantly, it directly upregulates Procr expression in the dividing cells, resulting in expansion of Procr+ progenitors around the wound.

## Discussion

In this study, we addressed the molecular mechanism in which OSE stem/progenitor cells sense the ovulatory rupture and promptly turns on proliferation and repair the wound. Our findings support the following model. Procr+ OSE progenitors have intrinsically lower level of Vgll4. Upon ovulatory rupture, decreased adherent junction at the proximity of rupture site promotes Yap nuclear localization. These intrinsic and extrinsic factors together lead to Yap signaling activation in Procr+ progenitors around the wound, which sequentially stimulates proliferation of the progenitors. Importantly, YAP activation directly upregulates Procr expression in the dividing cells, resulting in the expansion of Procr+ progenitors around the wound (Fig. 5m). Blocking Yap signaling in the progenitors by Yap-cKO or Vgll4-OE impairs the progenitors’ activities and hinders OSE repair. Furthermore, Procr function is essential for these progenitors. When Procr was deleted, stem cell property was lost hindering OSE repair.

YAP signaling promotes Procr+ cell expansion at rupture site through a combination of increased cell division and Procr expression. In the current study, YAP is particularly activated in Procr+ progenitor cells at rupture site. During the late stage of follicle development, the pre-ovulatory follicle forms a protrusion towards OSE. Subsequently, ovulation generates a rupture on OSE. These contiguous events might induce the thinning of OSE surrounding the pre- ovulatory follicles or at the proximity of the rupture site, leading to YAP activation. Another possibility is that the pre-ovulatory follicle protrusion or the release of oocytes induce special mechanical force on the OSE surrounding the wound, subsequently actives YAP signaling. Interestingly, at the rupture sites, Vgll4 is highly expressed in Procr- cells, preventing YAP pathway activation in those cells around the rupture sites. We showed here that reduced levels of Vgll4 in Procr+ progenitors likely contribute to the selective activation of YAP signaling in these cells. Further study should investigate what mechanism determines the lower expression of Vgll4 in Procr+ progenitor cells.

In the current study, we generated a new *tetO-Vgll4* mouse model that enables the overexpression of Vgll4 in specific cell type. The overexpression of Vgll4 in the progenitor of OSE has been validated using *Procr-rtTA* (Fig. 3g and Fig. S2d-e). The advantages brought by our *tetO-Vgll4* reporter will be of broad value in studies of Hippo-Yap signaling across all tissues.

Procr expression is initially found on the surface of vascular cells exerting an anti-coagulation role, by binding and activating protein C (PC) in the extracellular compartment^37^. More recently, studies from us and others have identified Procr as a stem cell surface marker in multiple tissues^7, 38–40^, but less is known regarding the function of Procr in stem/progenitor cells. In the current study, we demonstrate that, Procr is essential for the proliferation of Procr+ progenitor cells and OSE repair upon rupture. Our previous report indicated that PROCR concomitantly activates multiple pathways including ERK, PI3K-Akt-mTOR and RhoA-Rock-P38 signaling in breast cancer cells^41^. We speculate that similar intracellular pathways might be involved in the Procr+ OSE cells. Procr is regarded as a Wnt target gene from an *in vitro* screen in mammary stem cell culture^39^. In this study, we identify YAP as a novel upstream regulator of Procr. ChIP-qPCR and promoter luciferase experiments demonstrate that *Procr* transcription can be directly upregulated by YAP activation.

The phenomena of YAP promoting stem/progenitor cell expansion has been reported in various tissues ^30, 33, 34, 42–44^. Yet, in this process, less is known about how YAP maintains stem cell properties. To the best of our knowledge, this is the first report illustrating a mechanism through which YAP promotes cell proliferation, and simultaneously upregulates the expression of an essential stemness gene to maintain cell fate, leading to a rapid expansion of stem cell numbers around the wound. In summary, our study provides new evidence and molecular insights into how the OSE stem cell senses ovulatory rupture and promptly expands their numbers for repair. This may have a broad implication to understand the action of tissue stem cells during would healing in other tissues.

## ACKNOWLEDGMENTS

Thank to Dr. Chi Chung Hui of University of Toronto for helpful comments on the manuscript. This research was supported by grants from National Natural Science Foundation of China (31625020 to Y.A.Z. ; 32030025 and 31625017 to L. Z.); National Key Research and Development Program of China (2019YFA0802000 and 2020YFA0509002 to Y.A.Z.); the Chinese Academy of Sciences (XDA16020200 to Y.A.Z.); Fundamental Research Funds for the Central Universities (2020XZZX002-22 to J.F.); China Postdoctoral Science Foundation (2020TQ0260 to J.W.); Zhejiang Provincial Preferential Postdoctoral Foundation (ZJ2020150 to J.W.); Postdoctoral Exchange Fellowship Program 2021 (No.160 Document of OCPC, 2021) to J.W.; Shanghai Leading Talents Program to L. Z..

## AUTHOR CONTCONTRIBUTION

J.W. and Y.A.Z. designed the project and wrote the manuscript. J.W. performed most of the experiments including genetic crosses, FACS, RNA *in situ*, staining, cell culture, luciferase assay, qPCR and ovary phenotype analysis. L.H. and W.Y. performed ChIP-qPCR and provided YAP overexpression plasmids. Z.X. cloned the Procr promoter luciferase plasmids. L.B. and C.L. performed genetic crosses and superovulation. Y.L., Z.W. and Z.L. constructed the *TetO-Vgll4* mice and provided *YAP^fl/f^*^l^ mice. L.H, Z.L. and J.F. helped project design.

## Data Availability

All data generated or analysed during this study are included in the manuscript and supporting file; Source Data files have been provided for Figure 3-figure supplement 1-source data 1

N/A

## Ethics

Human Subjects: No Animal Subjects: Yes Ethics Statement: All mice were housed in the SIBCB animal facility under IVC standard with a 12-hr light/dark cycle at room temperature. Experimental procedures were approved by the Animal Care and Use Committee of Shanghai Institute of Biochemistry and Cell Biology, Chinese Academy of Sciences, with a project license number of IBCB0065.

## Methods

### Key Resource Table

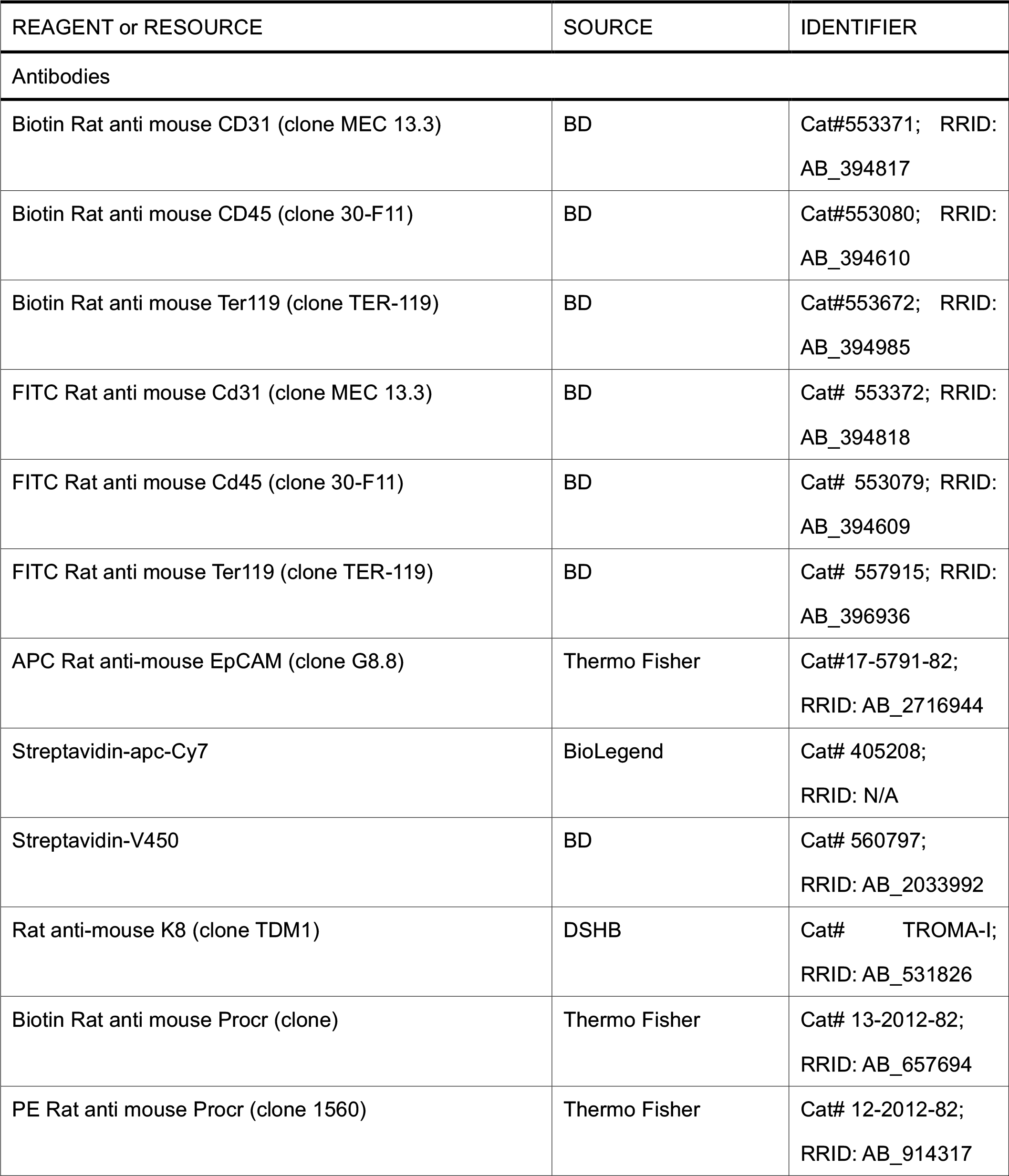
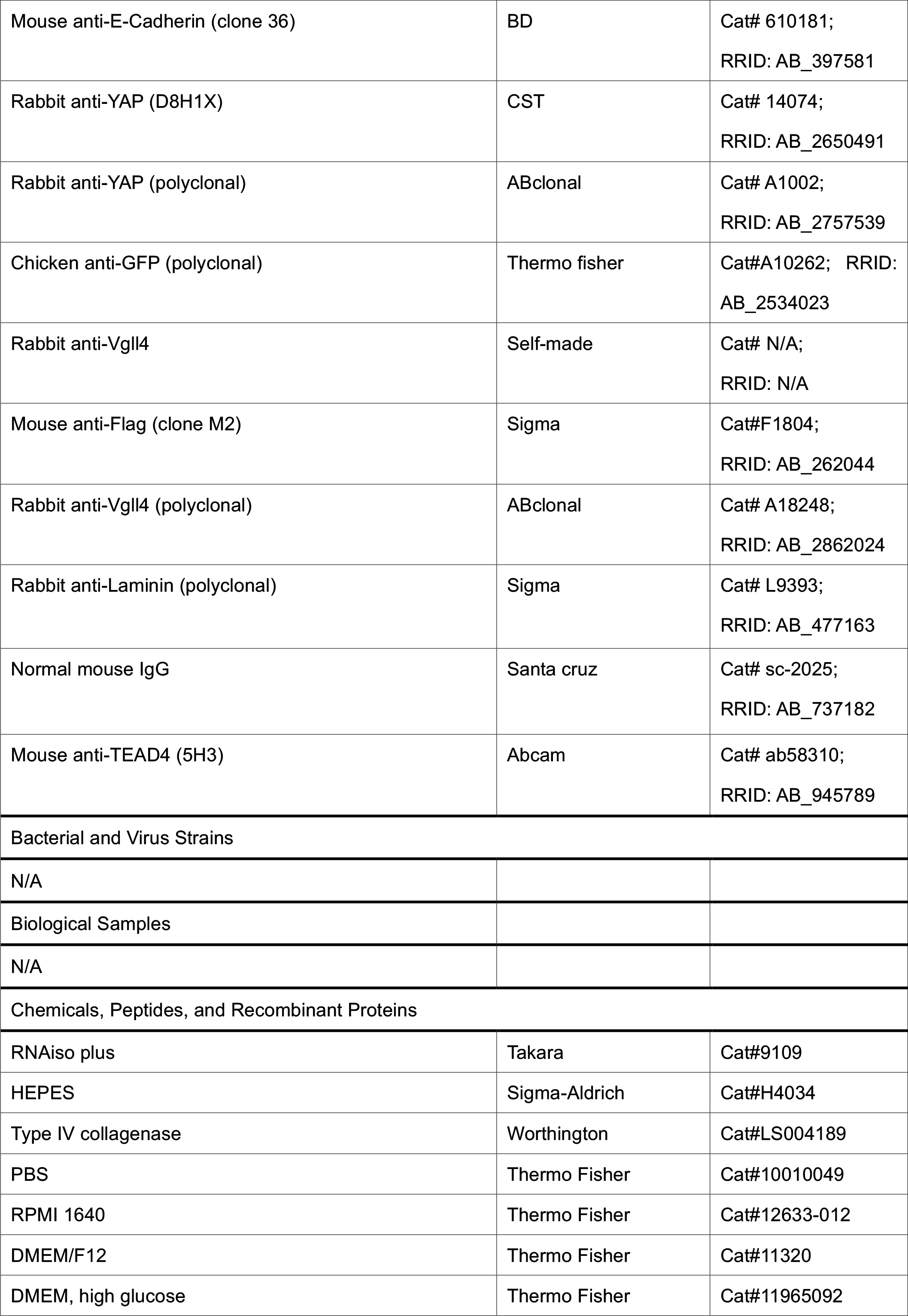
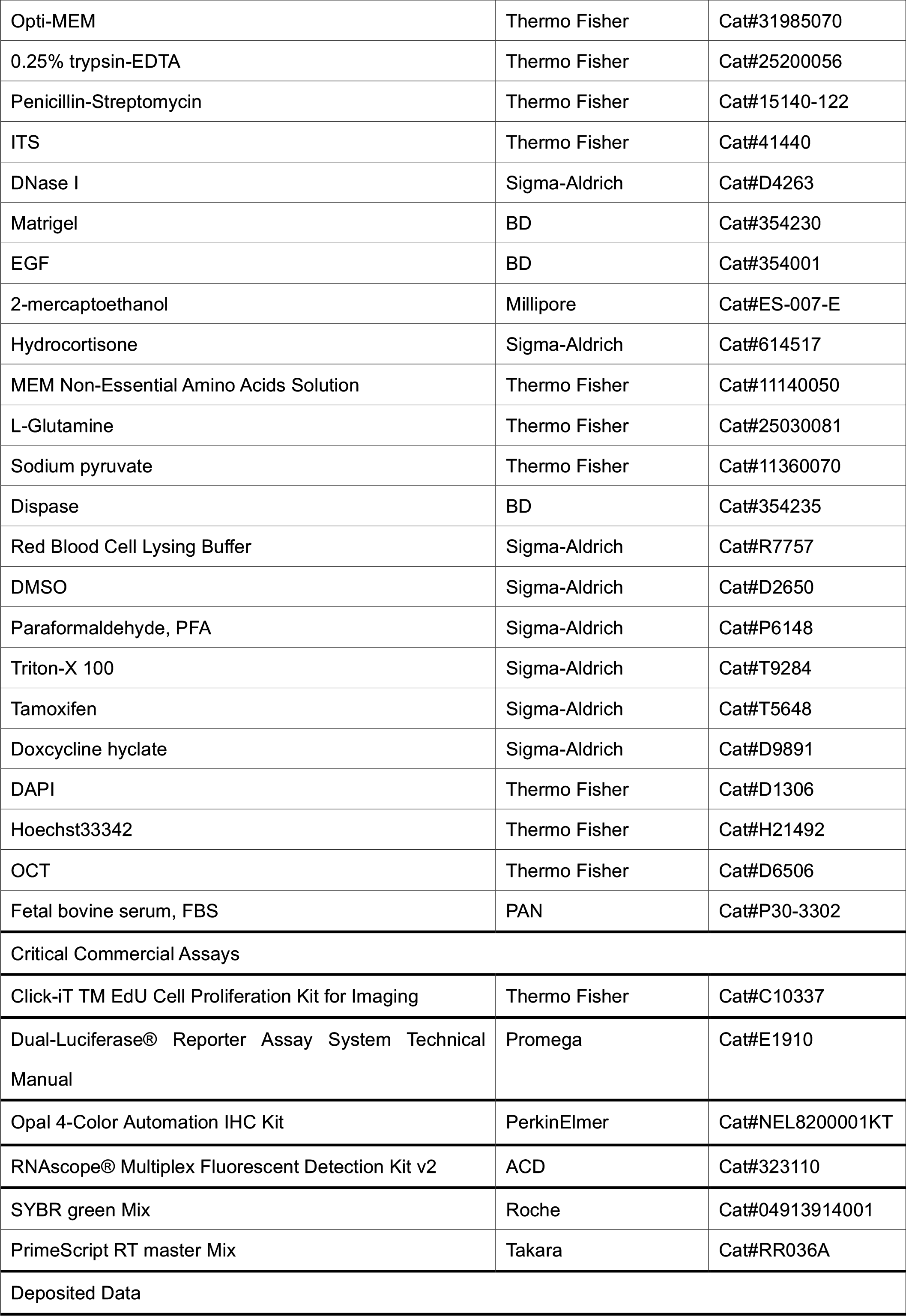
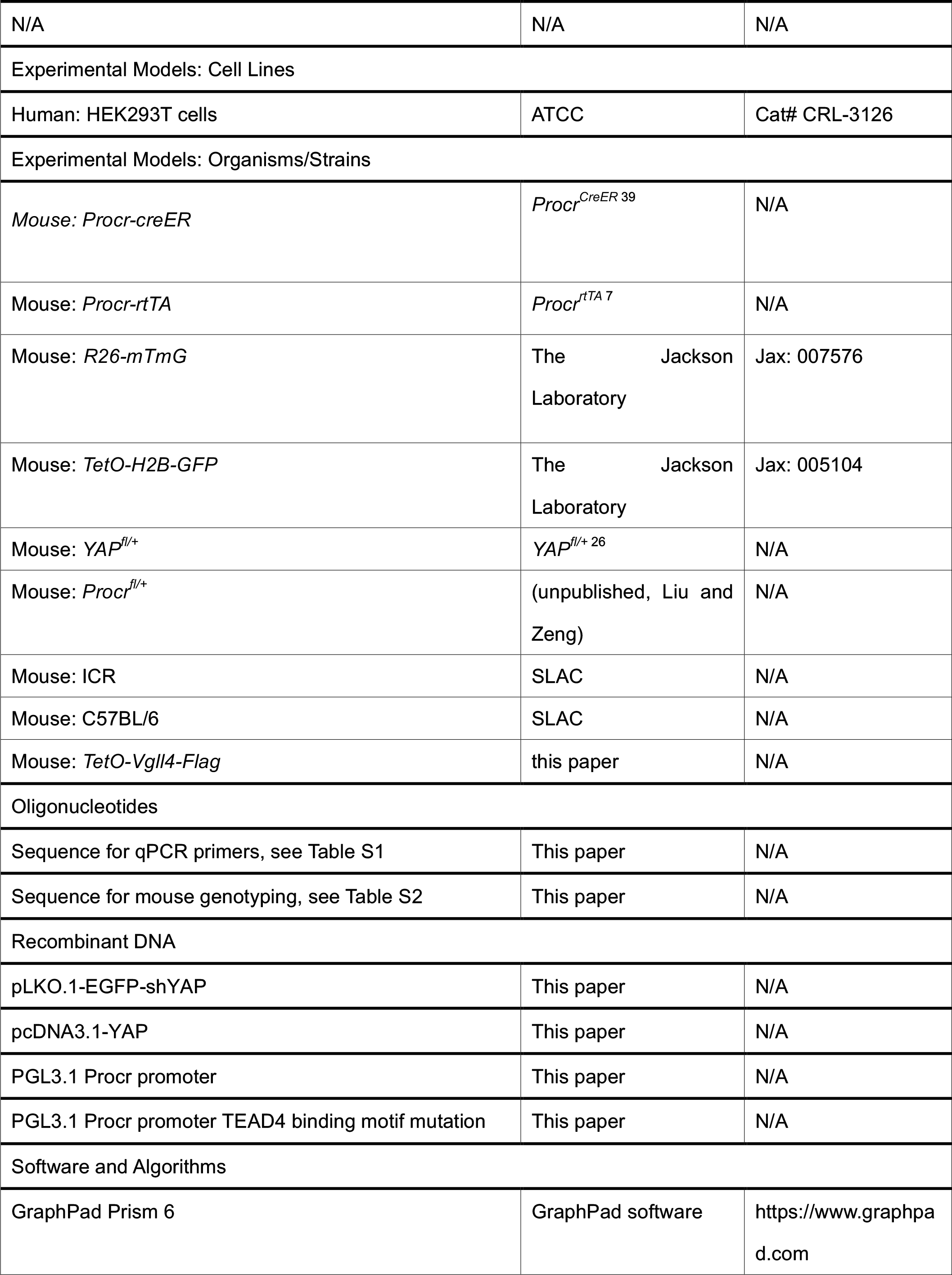
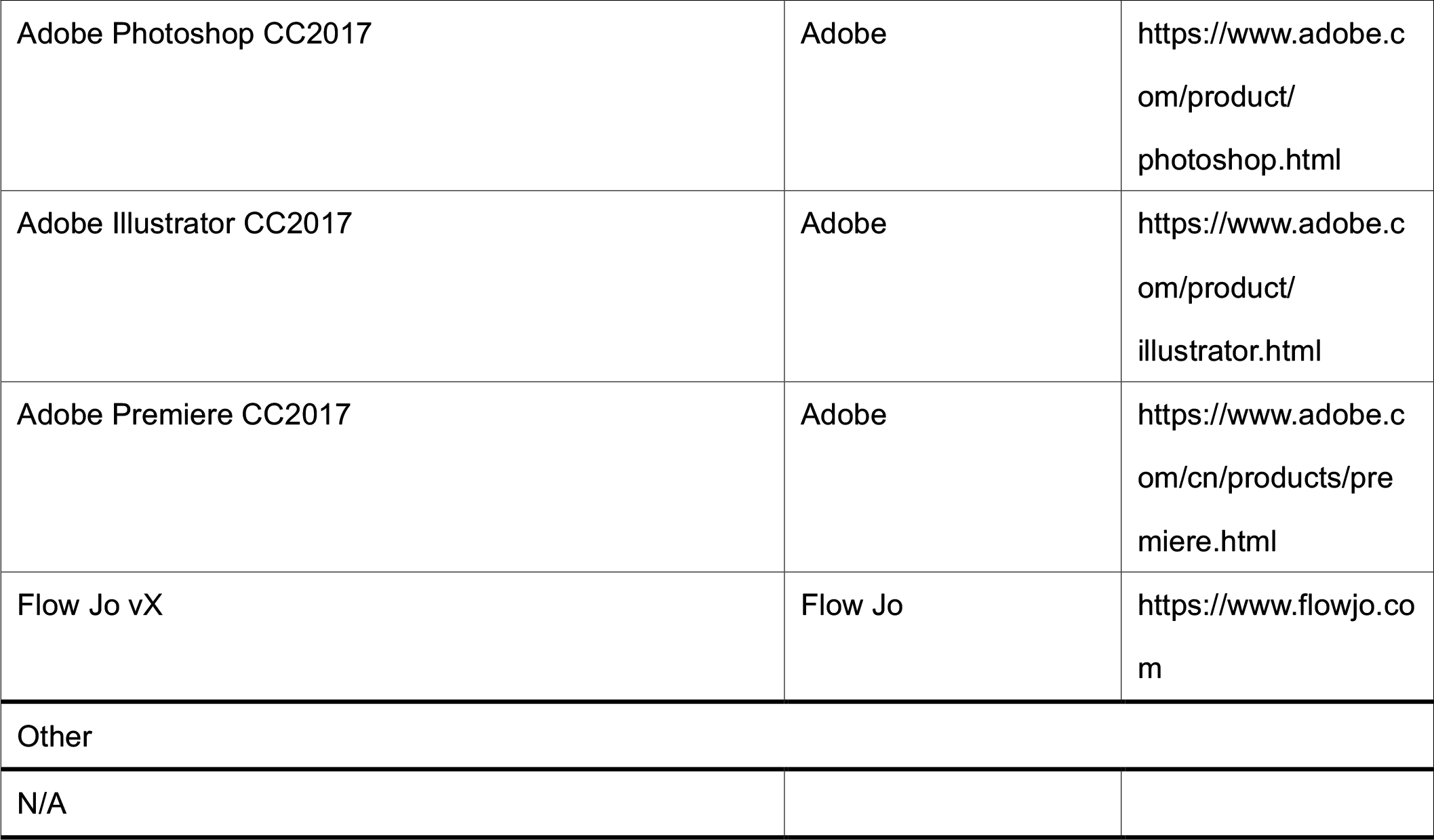

## LEAD CONTACT AND MATERIALS AVAILABILITY

Further information and requests for resources and reagents should be directed to and will be fulfilled by the Lead Contact, Yi Arial Zeng. All unique/stable reagents generated in this study are available from the Lead Contact with a completed Materials Transfer Agreement.

## EXPERIMENTAL MODEL AND SUBJECT DETAILS

### Experiment animals

*TetO-H2B-GFP^+/-^* (Stock: 005104), *R26-mTmG* (Stock: 007576) from Jackson Laboratories, *Procr^CreER^* ^39^, *Procr^rtTA^* ^7^, *YAP^fl/+^* ^26^, *Procr^fl/+^* (*unpublished,* Liu and Zeng), *TetO-Vgll4* were used in this study. The *TetO-Vgll4* mouse line was generated by knocking in a cassette of TetO-Vgll4-Flag-wpre-polyA behind 3’UTR of *Col1a1* gene (Fig. S2). All mice were housed in the SIBCB animal facility under IVC standard with a 12-hr light/dark cycle at room temperature. Both ovaries were used per mice and the number of mice per experiment was shown in figure legends. For targeted knockout *in vivo*, 4-5 weeks mice were administered with TAM diluted in sunflower oil by intraperitoneal (IP) injection at a concentration of 2mg per 25g body weight for two or three times (on every second day). For superovulation experiments, 4-5 weeks old mice were injected with 10 IU of pregnant mare serum gonadotropin (PMSG) by IP, followed by IP injection of 10IU of human chorionic gonadotropin (HCG) about 48 h later. For DOX feeding, doxycycline hyclate (DOX) was dissolved in drink water at a concentration of 1 mg/ml. Experimental procedures were approved by the Animal Care and Use Committee of Shanghai Institute of Biochemistry and Cell Biology, Chinese Academy of Sciences, with a project license number of IBCB0065.

### OSE cells isolation and flow cytometry

Ovaries from super-ovulated or 4-12 weeks old female mice were isolated, and the oviduct and bursa were carefully cleared out under dissect microscope. The ovaries were minced into pieces as small as possible, and then placed in 10 ml digest buffer (RPMI 1640with 5% fetal bovine serum, 1% penicillin– streptomycin, 25 mM HEPES and 300U/ml collagenase IV). After digestion at 37 °C, 100 rpm for about 1 hour, ovarian cells were obtained after centrifugation at 1000 rpm for 5 min. The red blood cells were lysed with buffer at room temperature for 5 min, and then single cells were obtained with 0.25% trypsin treatment at 37 °C for 5 min, followed by 0.1 mg/ml DNaseI incubation at 37 °C for 5 min with gently pipetting before filtering through 70 μm cell strainers. The single cells were incubated on the ice and in dark with the following antibodies at a dilution of 1:200: FITC conjugated, PE conjugated or biotinylated CD31, CD45, EpCAM-APC, Procr-PE, Procr-Biotin, Streptavidin-APC-Cy7, Streptavidin-V450. All analysis and sorting were performed using a FACSJazz (Becton Dickinson). The purity of sorted population was routinely checked and ensured to be > 95%.

### OSE cells 3D culture assay

FACS sorted OSE cells were resuspended with 60 μl 100% growth factor- reduced Matrigel and placed around the rim of a well of a 24-well plate, and allowed to solidify for at least 15 min at 37 °C in a 5% CO2 incubator before adding 0.5-1 ml culture medium. Colonies were grown for 7–9 days and the medium was changed every second days. The culture medium was prepared by adding 5% FBS, 4 mM L-glutamine, 1 mM sodium pyruvate, 10 ng/ml epidermal growth factor, 500 ng/ml hydrocortisone, 5 mg/ml insulin, 5 mg/ml transferrin, 5 ng/ml sodium selenite, 0.1 mM MEM non-essential amino acids, 10^−4^ M 2-mercaptoethanol into DMEM/F12. The organoid images were captured by Zeiss inverted microscope at day7-day9.

### Immunohistochemistry

For section staining, ovarian tissues were fixed in 4% PFA at room temperature for 15 min, following by washed with PBS for 3 times, dehydrated in 30% sucrose at 4°C overnight and embedded with Optimum Cutting Temperature (OCT). 16-18 μm tissue sections were incubated in 0.1% or 0.5% TritonX100 diluted with PBS (PBST) for 20 min and then 1 hour blocking using 10% FBS in PBST. Then sections were incubated with primary antibodies diluted in blocking buffer at 4°C overnight, followed by washes for 3 times (20 min per time). After wash, the sections were further incubated with secondary antibodies and DAPI diluted in blocking buffer for 2 hours at room temperature in dark, followed by washes for 3 times (20 min per time) and mounted with mounting medium.

For staining of cultured colonies, colonies were released from Matrigel by incubating with dispase for 20-30 min. Then the colonies were fixed in 4% PFA on ice for 10 min, followed by cytospin (Thermo Fisher) into slides and staining protocol described above.

For whole mouse ovary immunohistochemistry, mouse ovaries that cleared without bursa and oviduct were fixed with fresh 4% PFA at room temperature for 15 min in 4 ml Eppendorf tubes, followed by washing with PBST for three times (20 min per time). The staining of whole ovaries was then transferred into the 2 ml Falcon tubes using a dropper carefully. Ovaries were blocked for 1 hour using 10% FBS in PBST. Then, the ovaries were incubated with primary antibodies diluted in blocking buffer at 4 °C for 48 hours on a transference shaker with 10 rpm, followed by washing for three times (20 min per time) at room temperature. After washing, the ovaries were incubated with secondary antibodies diluted in blocking buffer for 24 hours at 4 °C in dark, and counterstained with DAPI. The ovaries could be stored in PBST at 4 °C for at least 2 weeks.

For YAP staining, Tyramide signal amplification assay (TSA staining) was used. Briefly, paraffin sections were rehydrated in histo-clear and gradual ethanol (100%,100%, 95%, 85%, 75%, 50%, 30%) and the TSA staining was performed using the Opal 4-Color Automation IHC Kit (PerkinElmer) following the manufacturer’s instructions. After TSA staining for YAP, staining for GFP and Krt8 was performed following protocol described above.

Tissue sections and organoids fluorescent images were captured using Leica DM6000 TCS/SP8 laser confocal scanning microscope with a 20×/0.75 or 40×/0.75 or 63×/0.75 IMM objective with 1-3 μm z-step. Confocal images were processed with maximum intensity projections.

Whole mouse ovarian fluorescent images were captured with inverted Leica TCS SP8 WLL at a 10×/0.75 objective, z-stack was ∼ 50–80 layers with 6-7 μm per layer, and the area was about 1.5 mm x 1.5 mm, which was about 1/6-1/4 of the adult ovary surface.

Primary antibodies used were: rat anti-Krt8 (1:250), rabbit anti-YAP (1:200), chicken anti-GFP (1:500), mouse anti-E-Cadherin (1:500), rabbit anti-Vgll4 (1:500), rabbit anti-Laminin (1:500). The secondary antibodies used were donkey anti-rat Cy3, donkey anti-rat Alexa Fluor 488, donkey anti-mouse Alexa Fluor 488, donkey anti-rabbit Alexa Fluor Cy3, donkey anti-mouse Alexa Fluor Cy3, donkey anti-rat Alexa Fluor Cy3, donkey anti-rat Alexa Fluor 647, donkey anti-mouse Alexa Fluor 647, donkey anti-rabbit Alexa Fluor 647, all secondary antibodies were used as 1:1000. At least three times repeats were done per tissue block. Only representative images were shown.

### Western Blotting

Digested cells were lysed in SDS-PAGE loading buffer and boiled for 10min. Proteins were separated by SDS-PAGE and transferred to nitrocellulose membrane (GE company). Bolts were blocked with 3% BSA in TBST (50 mM Tris-HCl, 150 mM NaCl, 0.05% Tween-20, pH 7.5) for 1 hour and incubated with primary antibodies at 4 °C overnight, followed by incubated with secondary IgG-HRP antibodies for 2 hours at room temperature. Protein bands were visualized with chemiluminescent reagent and exposed to Mini Chemiluminescent Imager.

### RNA *in situ*

In situ hybridization was performed using the RNA scope kit (Advanced Cell Diagnostics) following the manufacturer’s instructions. *Pro*cr probes (REF#410321) and *Cyr61* probes (REF#429001) were ordered from Advanced Cell Diagnostics. After in situ hybridization, TSA method was used for Krt8 staining following the using the Opal 4-Color Automation IHC Kit (PerkinElmer) following the manufacturer’s instructions. The images were captured using Leica DM6000 TCS/SP8 laser confocal scanning microscope with a 63×/0.75 IMM objective.

### EdU labelling assays

The proliferation of OSE cells *in vivo* or *in vitro* was measured by 5- ethynyl-29- deoxyuridine (EdU) uptake. Briefly, mice were injected with 100 μl EdU (2.5 mg/ml in dimethyl sulfoxide) for 12 h. Then ovaries were harvested for section, following by EdU color staining using Click-iT EdU Alexa Fluor Imaging Kit (prepared according to the manufacturer’s instructions). After washed with PBS for 3 times (10 min per time), EdU color development was performed following manufacturer’s protocol. After EdU signal developing, sections were blocked in blocking buffer for 1 h at room temperature followed by antibody staining and mounted with mounting medium for imaging and quantification.

### Living image of cultured OSE cells

OSE cells were isolated from the mice and cultured on glass for 3-4 days. DOX was added into the medium 1 day and Hoechst33342 was added 30 mins before image. Live-cell imaging was performed at 37 °C on a Zeiss Celldiscoverer 7 with perfect focus system. Cells were imaged at 1 time per 5 mins for 24 hours with 70% laser power.

### Chromatin Immunoprecipitation-qPCR (ChIP-qPCR)

Cultured primary OSE cells were crosslinked in a final concentration of 1% formaldehyde (Sigma) PBS buffer for 15 min at 37 °C, then added glycine to stop crosslinking. Chromatin from nuclei was sheared to 200–600 bp fragments using ultrasonic apparatus, then immunoprecipitated with antibody of TEAD4 (ab58310, Abcam) or normal mouse IgG (sc-2025, Santa Cruz) overnight. Antibody/antigen complexes were recovered with Protein A/G PLUS-Agarose (sc-2003, Santa Cruz Biotechnology) for 2 hours at 4 °C. After washing, the chromatin was eluted, de-crosslinked and digested. The immunoprecipitated DNA was collected with QIAQIUCK PCR Purification Kit (QIAGEN)). Purified DNA was performed with ChIP-qPCR. Assessing the enrichment of the proteins of interest on the targeting region by calculating the value of “fold over IgG”. ChIP-qPCR primers used were as follows.

Ctfg CHIP-F, TGTGCCAGCTTTTTCAGACG;

Ctfg CHIP-R, GAACTGAATGGAGTCCTACACA;

Procr CHIP-F, ATATCCGAGCTACACACGGC;

Procr CHIP-R, GTGAATGCACACACACACCC;

Negative Ctrl CHIP-F, GATCAACGCAGGGGAGAGAG;

Negative Ctrl CHIP-R, TATCCCCACTGCCCAGAAGA.

### Preparation of Procr promoter luciferase reporter and luciferase assay

The DNA sequence of Procr promoter containing TEAD4 binding sites (about 2kb before the initiation codon) were amplified by PCR, separated by agarose gel, purified by Gel Extraction Kit, and then cloned into pGL3-promoter vector. Luciferase assays were performed in 293T cells with the pGL3-Procr promoter luciferase reporter described above 0.2 mg reporter plasmid were transfected together with CMV-Renilla (0.005 mg) to normalize for transfection efficiency. For luciferase assays in overexpression plasmid-transfected cells, cells were transfected with the indicated plasmids and reporter plasmid together, and then the luciferase activity was measured 36 hours later using Dual-Luciferase® Reporter Assay System Technical Manual kit following manufacturer’s protocol.

### Cell Culture, viral production and infection

HEK293T, C3H10 were obtained from American Type Culture Collection (ATCC) and cultured in Dulbecco modified essential medium (DMEM) supplemented with 10% FBS plus 1% penicillin and streptomycin antibiotics at 37 °C in 5% CO2 (v/v). For cells cultured on different modulus of elasticity, hydrogel substrates with tunable mechanical properties were prepared following the previous protocol ^45^, and the glass was as solid control. HEK 293T cells were used to produce lentivirus. When cells were up to 80-90%, indicated constructs and packaging plasmids transfection was performed in Opti-MEM, and the media were replaced 12 hours later. Viral supernatants were collected 48-72 hours after medium change and filtered through a 0.45 μm filter, followed by concentration. For primary OSE cells infection, concentrated virus was diluted in the culture medium along with 1:100 polybrene.

### RNA isolation and quantitative real-time PCR

Total RNA was isolated from fresh OSE cells or cultured cells lysed with Trizol according to the manufacturer’s instructions. The cDNA was generated from equal amounts of RNA using the SuperScriptIII kit. qPCR was performed on a StepOne Plus (Applied Biosystems) with Power SYBR Green PCR Master Mix. RNA level was normalized to *Gapdh*. The cycling condition was as 10 min at 95°C for initial denaturing, 40 cycles of 15 s at 95°C for denaturing, 1 min at 60°C for annealing and extension, following by melt curve test. Primers used were as follows.

*Procr*-F, CTCTCTGGGAAAACTCCTGACA;

*Procr*-R, CAGGGAGCAGCTAACAGTGA;

*Vgll4*-F, ATGAACAACAATATCGGCGTTCT;

*Vgll4*-R, GGGCTCCATGCTGAATTTCC;

*YAP*-F, GCCATGCTTTCGCAACTGAA;

*YAP*-R, CAAAACGAGGGTCCAGCCTT;

*Cyr61*-F, TCGCAATTGGAAAAGGCAGC;

*Cyr61*-R, CCAAGACGTGGTCTGAACGA.

### Quantification and statistical analysis

For quantification of YAP+, Vgll4+ and EdU+ cells, 40 OSE cells at the both edges of ruptured sites (20 OSE cells at one side of rupture site) was identified as rupture regions, while other regions as non-rupture regions. At least 30 rupture regions and 30 non-rupture regions were counted. For quantification of the diameter of rupture, the longest diameter was counted, and at least 20 rupture sites were counted. For quantification of mG+ clone sizes, about 0.3 mm^2^ circle centered on ruptured sites was identified as rupture regions. At least 30 rupture regions were counted. For quantification of colonies size, diameters of the colonies were measured using Zeiss software.

Statistical analyses were calculated in GraphPad Prism (Student’s t-test or One-way ANOVA). For all experiments with error bars, the standard error of measurement (s.e.m) was calculated to indicate the variation within each experiment. All the p values were calculated using GraphPad PRISM 6 with the following significance: n.s. p > 0.05; * p < 0.05; ** p < 0.01; *** p < 0.001. Statistical details for each experiment can be found in the figures and the legends.

---

**Fig S1.**
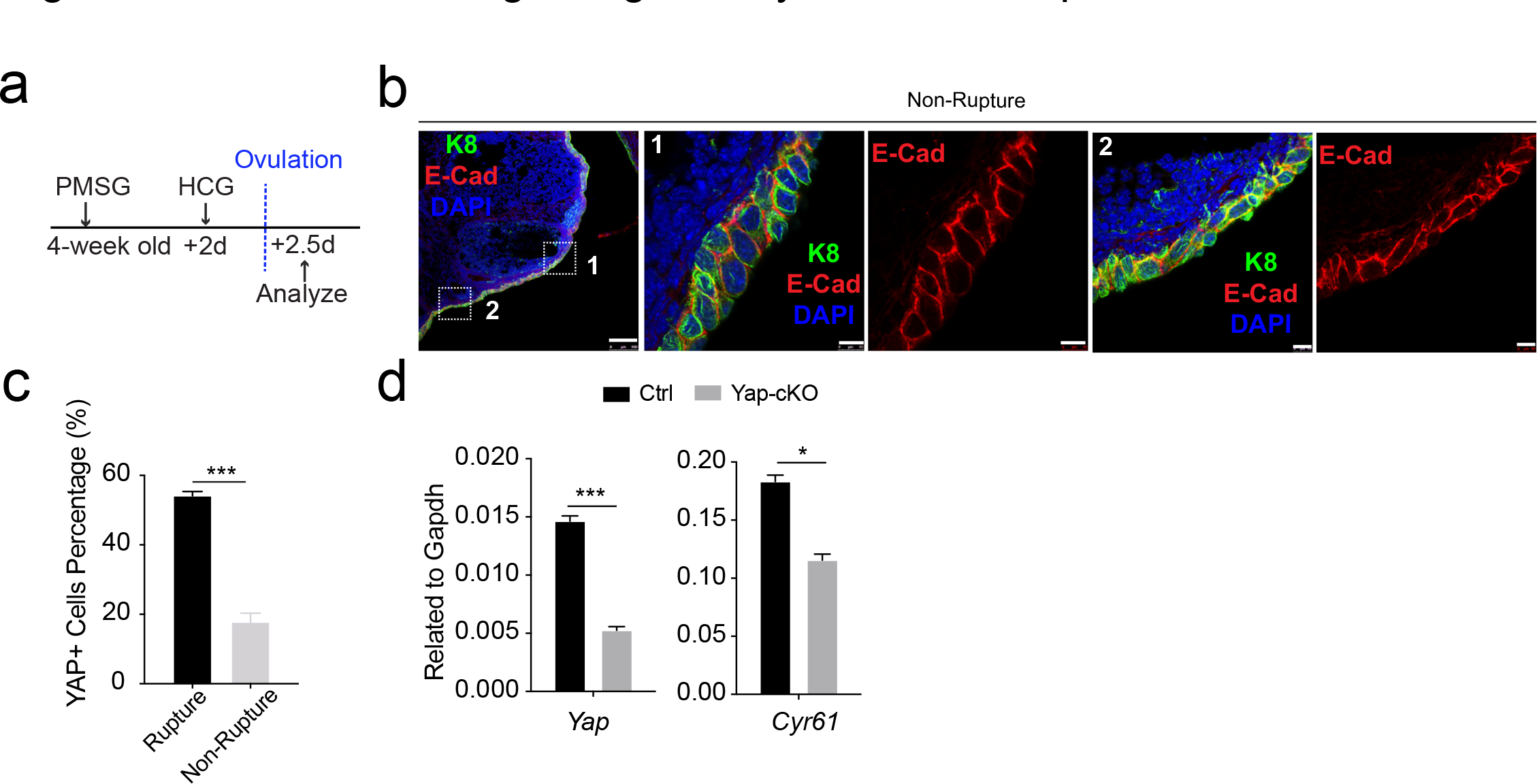
Increased YAP signaling activity at OSE of rupture sites. a, Illustration of superovulation strategy. 4-week old mice were administrated with PMSG, following by HCG 2 days later. The ovaries were harvested 0.5 day after HCG injection (ovulation). b, Confocal images showed abundant E-cad expression in the OSE of non- rupture sites. Scale bar, 100μm for zoom out and 10μm for zoom in. n=3 mice and more than 15 images. c, Quantification of the percentage of OSE cells with YAP nuclear localization at the rupture sites and non-rupture sites. n=3 mice. Unpaired two-tailed t test is used for comparison. ***P < 0.001. d, qPCR analysis validated the deletion efficiency of *Yap* and the downregulation of the expression of YAP target *Cyr61* in total OSE cells of Yap- cKO mice compared with Ctrl mice. Data are pooled from three independent experiments and displayed as mean±s.e.m. Unpaired two-tailed t test is used for comparison. ***P < 0.001, *P < 0.05.

**Fig S2.**
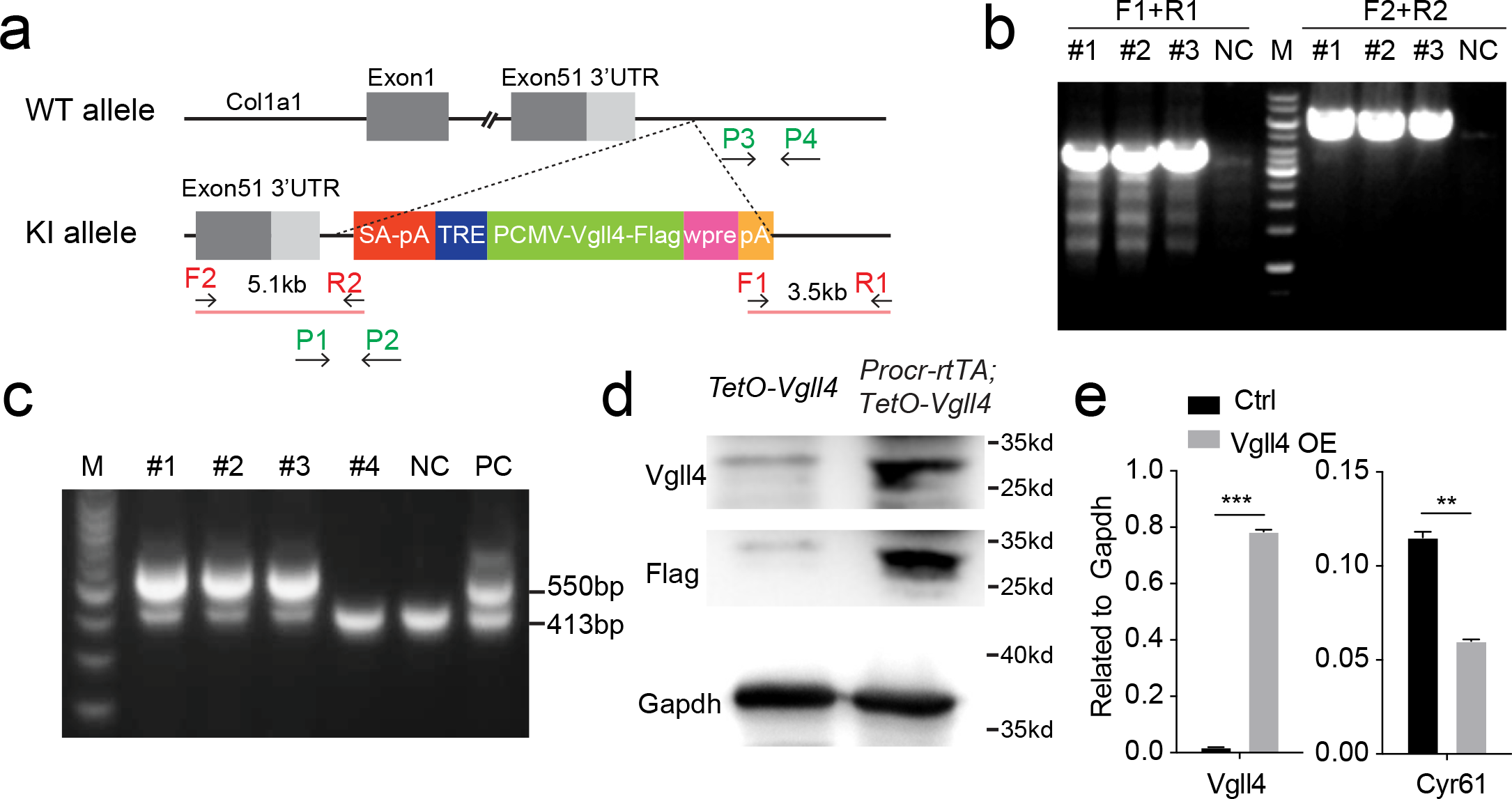
Construction of *TetO-Vgll4* mouse model. a, Targeting strategy for the generation of *TetO-Vgll4* knock-in mouse. Designs of ES clone genotyping primers (red) and mouse genotyping primers (green) are as indicated. b, ES clone genotyping PCR indicating three successful knock-in (KI) clones. NC, negative control with no DNA input. c, Genotyping PCR results indicate pup #1,2,3 is heterozygote, #4 are wildtypes. A wild-type mouse was used as negative control (NC) and a positive ES clone was used as positive control (PC). d, Western blotting validated the overexpression of Flag and Vgll4 in the cells of Vgll4-OE mice compared with Ctrl mice. One of 3 independent experiments is shown. e, qPCR analysis validated the overexpression of *Vgll4* and downregulation of *Cyr61* in total OSE cells of Vgll4 OE mice compared with Ctrl mice (d). Data are pooled from three independent experiments and displayed as mean±s.e.m. Unpaired two-tailed t test is used for comparison. ***P < 0.001, **P < 0.01.

**Fig S3.**
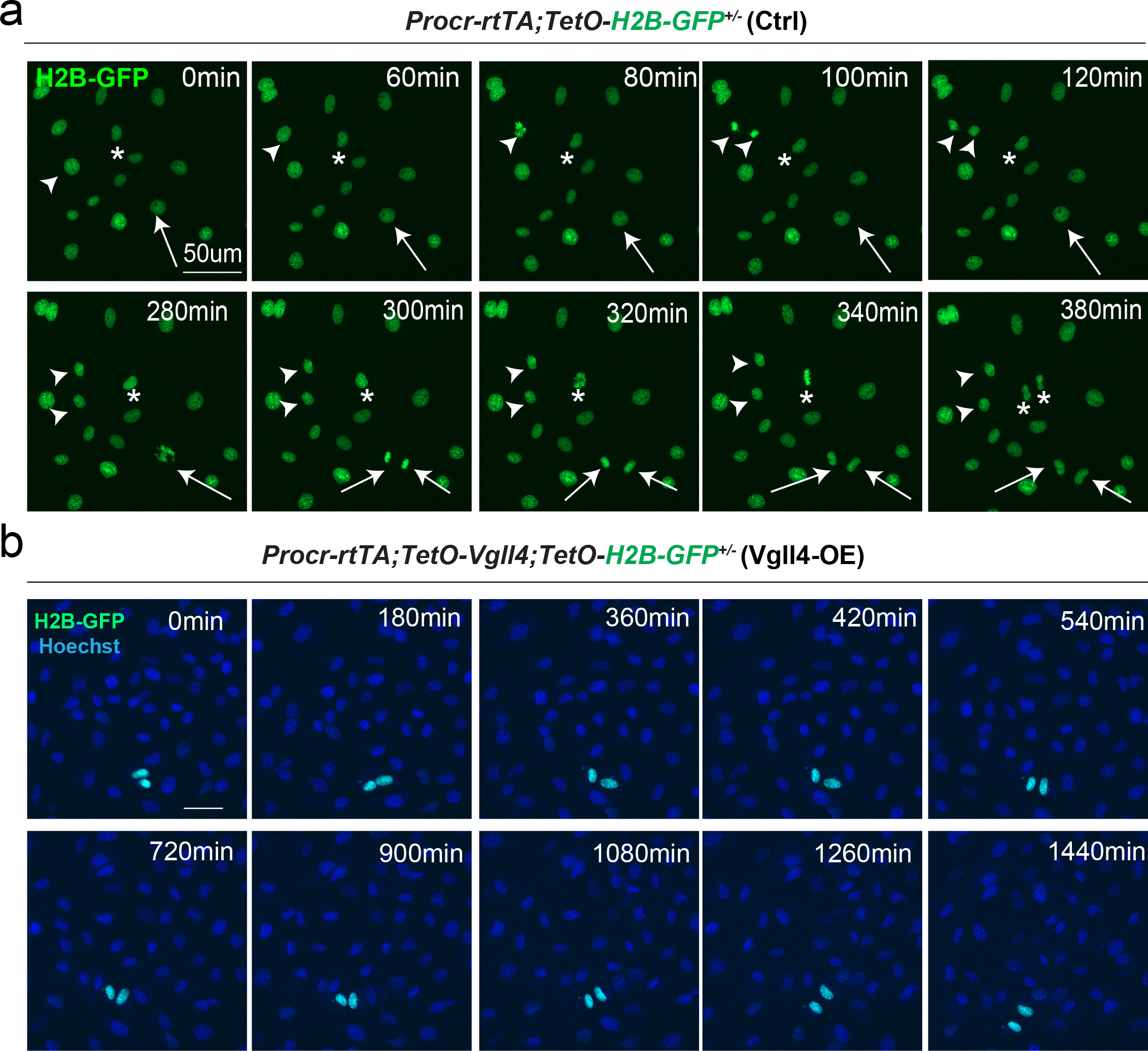
YAP promotes Procr+ cells expansion. a-b , OSE cells were isolated from *Procr-rtTA;TetO-H2B-GFP^+/-^* (Ctrl) or *Procr- rtTA;TetO-Vgll4^+/-^;TetO-H2B-GFP^+/-^* (Vgll4-OE) mice and cultured on the glass. In control, almost all cells are H2B-GFP+ (Procr+) in such stiff culture condition (a). Living images for 6 hours showed many cases of H2B-GFP+ (Procr+) cells (a) (examples in *, arrow, arrowhead in a). In Vgll4-OE cells, there were drastically less H2B-GFP+ (Procr+) cells, and living imaging of 24 hours showed no incidence of cell division (b). Scale bar, 50μm. n=at least 3 views.

**Fig S4.**
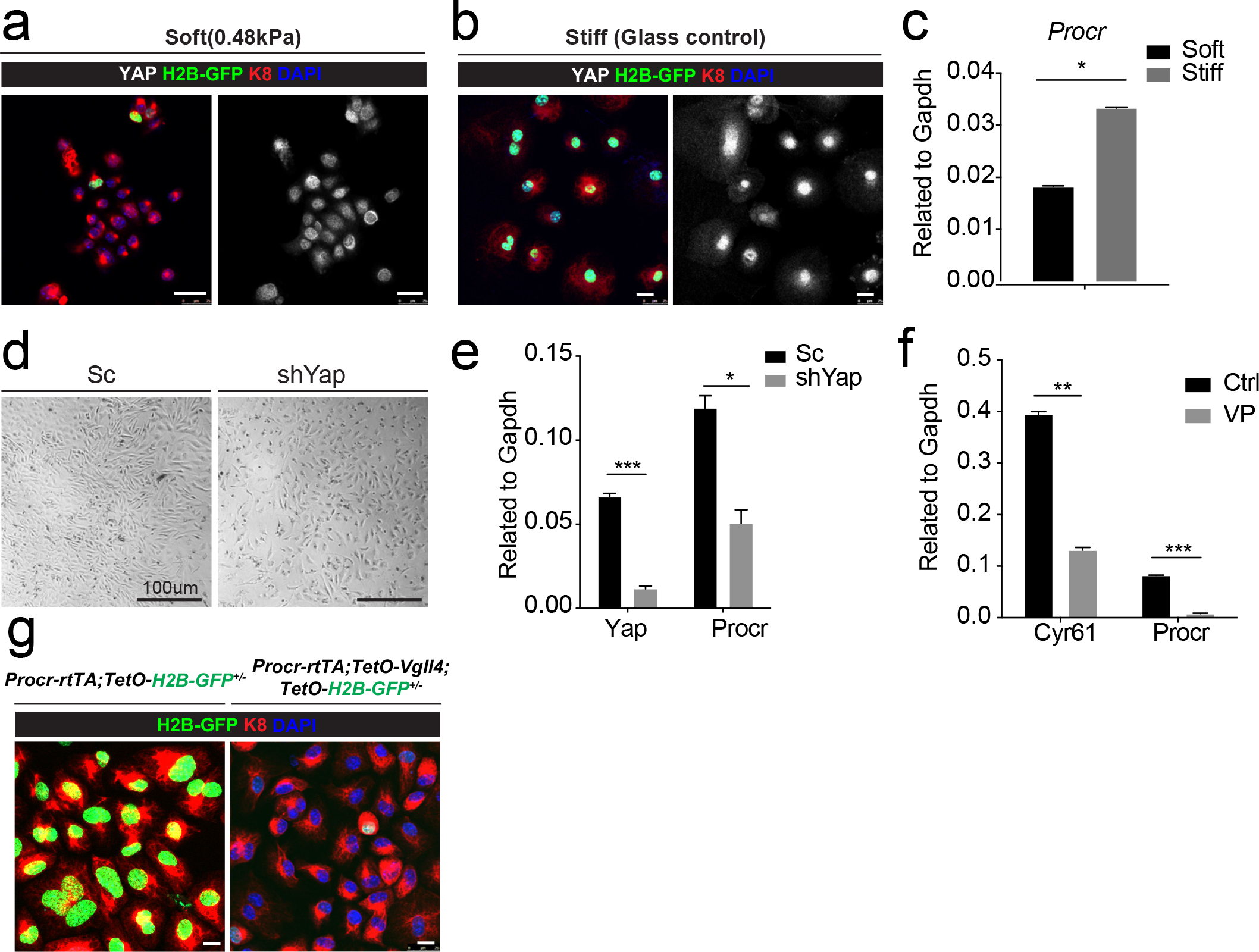
YAP induces Procr expression. a-c , OSE cells were isolated from *Procr-rtTA;TetO-H2B-GFP^+/-^* mice and cultured upon the soft hydrogel (a) or glass (b). Confocal images showed more H2B-GFP+ cells upon glass compared with soft hydrogel (a-b). Scale bar, 20μm, n=15 images. qPCR indicated Procr expression was upregulated upon glass culture (c). d-e, OSE cells isolated from wildtype mice were infected with Scramble (Sc) or Yap shRNA (shYap) virus. and then cultured on glass. OSE cells were harvested on culture day 4 (d). qPCR showed knockdown of Yap repressed *Procr* expression (e). Scale bar, 100μm, n=15 images. f, OSE cells were isolated from wildtype mice and cultured on glass. Verteporfin (VP) was added into the medium before harvest. qPCR showed that VP treatment inhibits *Cyr61* and *Procr* expression. g, OSE cells were isolated from *Procr-rtTA;TetO-H2B-GFP^+/-^* (Ctrl) and *Procr-rtTA;TetO-Vgll4^+/-^;TetO-H2B-GFP^+/-^* (Vgll4-OE) mice and cultured on the glass. Confocal images showed dimmer H2B-GFP expression in Vgll4-OE compared to Ctrl. Scale bar, 10μm, n=15 images. For all qPCR results, data are pooled from 3 independent experiments and presented as mean±s.e.m. Unpaired two-tailed t test is used for comparison. ***P<0.001, **P<0.01, *P<0.05.

